# PD-1 checkpoint blockade activates germinal center follicular T cell programs that disrupt type 2 isotype-specific antibody homeostasis

**DOI:** 10.1101/2021.09.27.462076

**Authors:** Andrew G. Shuparski, Brett W. Higgins, Karen B. Miller, Louise J. McHeyzer-Williams, Michael G. McHeyzer-Williams

## Abstract

Multiple CD4 T cell dependent tolerance mechanisms control adaptive B cell immunity to environmental antigens. We recently demonstrated a PD-1 checkpoint within steady-state splenic germinal centers (GC) that constrains the maturation of type 2 IgG1 isotype-specific antibody homeostasis. Here, we utilized single cell-indexed custom RNA-sequencing to probe the follicular T cell mechanisms directly targeted by acute PD-1 blockade. We find a pre-existing subset of follicular helper T (T_FH_) cells that express type 2 immune response properties (T_FH_2) with exaggerated pathways of TCR activation, cytokine signaling, and enhanced cell-cell contact upon acute PD-1 blockade. This selective amplification of the T_FH_2 program significantly increases predicted molecular connections to type 2 IgG1 GC B cells that dominate limited changes in GC localized follicular regulatory T (GC T_FR_) cell programs. These studies demonstrate how type 2 isotype-specific adaptive B cell tolerance is selectively disrupted by acute PD-1 blockade to reveal the modular regulatory mechanisms that control splenic GC dynamics at homeostasis.

**One Sentence Summary:** Acute PD-1 blockade alters the regulatory dynamic of the steady state germinal center to drive the maturation of IgG1 GC B cells towards PC differentiation in a process mediated by type 2 like TFH effector molecules.

**HIGHLIGHTS:**

- Acute PD-1 blockade enhances the steady state splenic TFH program
- PD-1 blockade selectively exaggerates a Type 2 like TFH module
- GC TFR cells are minimally impacted by blockade
- PD-1 restrains predicted TFH2 functional contacts with IgG1 GC B Cells

## INTRODUCTION

Homeostatic or “steady state” immunity must be finely tuned to respond to pathogens while enforcing tolerance against commensal bacteria ^1,2,34^. Enabling this balance is a multifactorial process that also requires tight regulation and control over adaptive B cell immunity provided by specialized CD4 T cell follicular compartments ^5, 6^. These follicular T cell compartments can be divided into helper (T_FH_) and regulatory (T_FR_) subsets that interact in dynamic opposition to regulate GC B cell function and fate ^7, 8^. Programmed cell death protein 1 (PD-1) is expressed at high levels by both follicular T cell subsets ^9,10,11^ indicating their capacity for GC B cell checkpoint control. It remains unclear what molecular mechanisms control these cellular interactions and how they impact adaptive B cell tolerance at homeostasis.

PD-1 plays a prominent role in inducing T cell exhaustion in cancer and chronic viral anergy models ^12^. Blockade of PD-1 signaling in these contexts can improve T cell function ^13,14,15,16^ and this observation that has been extended to the clinic with the FDA approved immunotherapies Nivolumab (Bristol-Myers Squibb, USA) and Pembrolizumab (Merk, USA). Interference with PD-1 signaling has led to either a decrease ^17,18,19^ or augmentation of humoral immunity ^14, 20,21,22^. Here, we focus on the role of PD-1 in regulating adaptive B cell tolerance. We recently demonstrated that acute PD-1 blockade releases type 2 IgG1 GC B cell maturation to produce massive local expansion and differentiation of IgG1 plasma cells (PC) and antibodies ^23^. Released IgG1 specificities target pre-existing foreign antigens on gut microbiota and impact local microbial composition. Surprisingly, there was no evidence for autoimmunity across multiple immunization formats and timepoints, and no overt adjuvant action in a vaccine context. We propose that this adaptive tolerance mechanism acts directly through GC-localized follicular CD4 T cell compartments that selectively constrains type 2 isotype-specific GC B cells to restrict antibody affinity maturation and IgG1 PC differentiation.

CD4 T cell compartments can be broadly organized across three major classes of immune response. Divergent lineage defining protein and transcriptional signatures assort with type 1 inflammatory (targeting intracellular microbes), type 2 anti-inflammatory (targeting extracellular microbes), and type 3 mucosal (protecting barrier surfaces) immune response modules ^24,25,26,27^. Conventional non-T_FH_ effector (ETH) and regulatory T (T_reg_) cells are typically considered under this conceptual framework but more recently this division has been proposed for follicular T cell compartments ^28,29,30,31,32^. In this new schematic, T_FH_ cells segregate according to the antibody isotype they promote in activated B cells and subsequently control in an isotype-specific manner throughout primary and recall immunity.

Selective Tbet dependency of type 1 inflammatory IgG2a B cell memory ^33^ suggests type1 response modules of T_FH_ cells may selectively control these outcomes. In contrast, murine type 2 IgG1 antibodies bind inhibitory Fc receptors ^34^ that can resolve inflammation in the context of autoimmunity ^35^. The more inflammatory IgE antibody isotype, produced downstream of IgG1, is also considered a type 2 response with recent evidence for a separable T_FH_13 compartment controlling this pathway ^36^. Here, we probe the follicular T cell mechanisms of homeostatic control for the type 2 IgG1 isotype-specific pathway that is released upon acute PD-1 blockade ^23^.

We use multiplexed protein and transcriptional single cell strategies in a Foxp3 reporter model to interrogate both arms of the murine splenic T follicular cell response following acute PD- 1 blockade. We find a subset of TFH cells exhibiting type 2 properties and markers associated with localization in the GC that responds markedly to PD-1 blockade. This “T_FH_2” cell subset upregulates an expansive transcriptional program including key genes affiliated with cytokine signaling, cell to cell contact, and TCR activation. Following PD-1 blockade T_FH_2 cells but not T_FH_1 or T_FR_ cells increase their potential to provide help to cycling IgG1 GC B cells. Amplification of the T_FH_2 program further alters the local regulatory balance to disrupt anti-inflammatory GC B cell homeostasis. We reveal mechanisms of T follicular cell control of germinal center dynamics and insight into the emergent properties of isotype-specific regulation ^23^ and its particular role in maintaining adaptive B cell tolerance.

## RESULTS

### Divergent programs of CD4 T cell subsets in the steady-state spleen

The steady-state spleen contains pre-existing GC B cells, memory B cells and plasma cells across all CD4-dependent class-switched antibody isotypes ^23^. Two fractions of non-T_FH_ effector (ETH) cells comprise approximately half of the pre-existing ’activated’ T cell compartment (**Fig. 1a**). Foxp3 expression clearly delineates the conventional T regulatory (T_reg_) fraction while CD25 expression further segregates two subsets of Cxcr5 expressing Foxp3^+^ T_FR_. The CD25^hi^ T_FR_ are thought to regulate early events at the T-B border ^8^ while CD25^lo^ T_FR_ express higher levels of PD-1 and to localize to the GC ^11, 37, 38^. While low in numbers, Cxcr5^hi^ PD-1 expressing T_FH_ comprise a significant fraction of the spleen at homeostasis. For gene expression analysis, we required a Foxp3^Thy1.1^ reporter model to access and segregate these rare ’activated’ and regulatory CD4 T cell compartments without fixation (**Fig-1b**).

**Figure 1.**
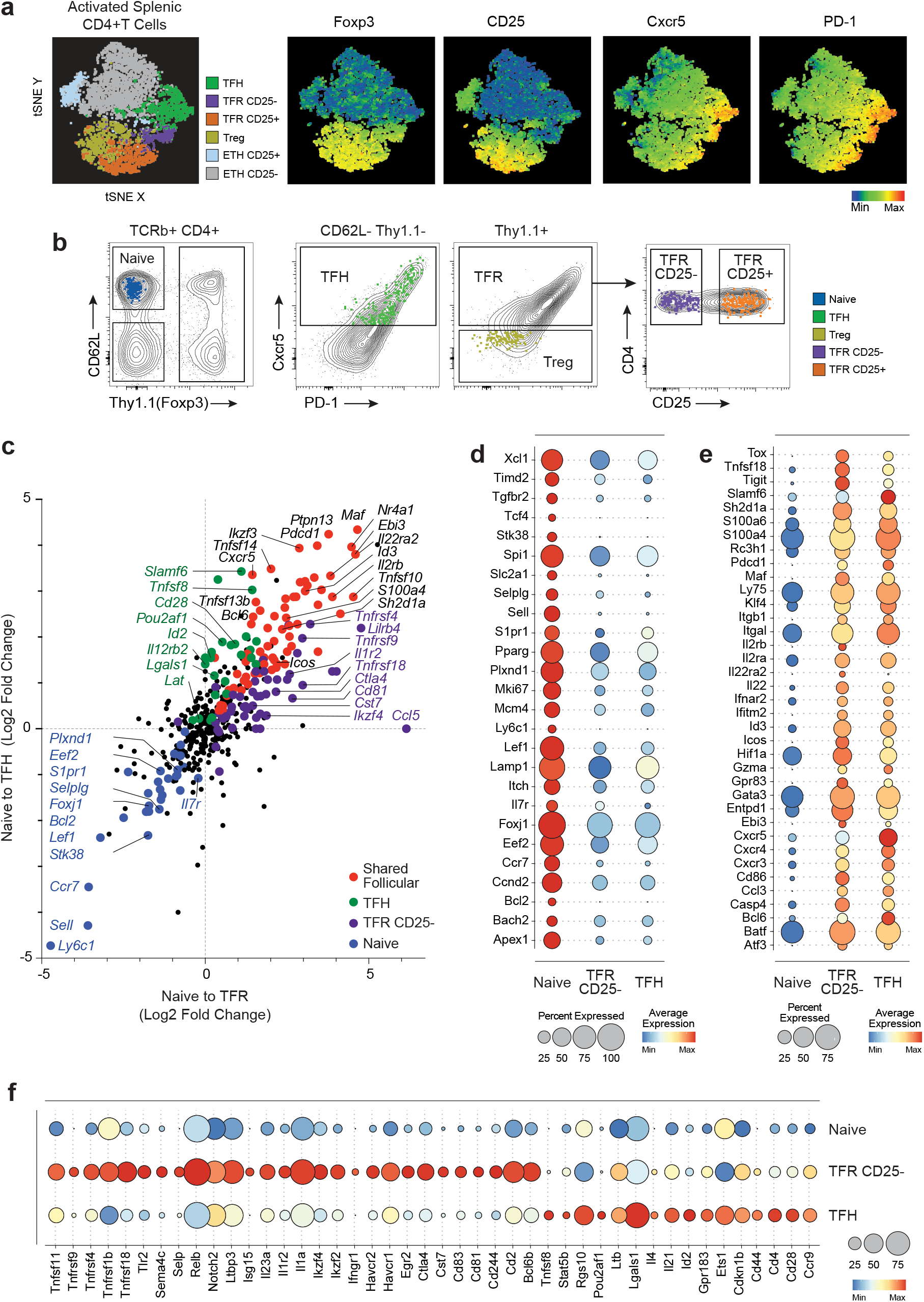
Divergent programs of CD4 T cell subsets in the steady state spleen. **a)** Foxp3^Thy1.1^ reporter mice t-distributed stochastic neighbor embedding (tSNE) flow cytometry analysis of splenic CD4 T cells (B220-CD8-, CD4+TCRb+, CD62L-CD44+ and/or Foxp3+). Six combined parameters (Foxp3, CD25, Cxcr5, PD-1, Cxcr3 and ICOS) were used to cluster the cells (left panel). T follicular helper (TFH), T follicular regulatory (TFR), T regulatory (Treg) and T effector helper (ETH) cells are colored as indicated (n=8000 cells). Distribution of individual proteins Foxp3, CD25, Cxcr5 and PD-1 are overlayed (right) from minimum (blue) to maximum (red) intensity for each fluorophore. **b)** Flow cytometric analysis of TCRb+ CD4+ T cells from steady state mice with overlaid gated index sorted cells. Naïve (Thy1.1- CD44- CD62L+) n=200 colored blue, TFH (Thy1.1- CD44+ CD62L- Cxcr5+) n=363 green, Treg (Thy1.1+ Cxcr5-) n=65 yellow, TFR (Thy1.1+ Cxcr5+) either CD25- n=363 purple, or CD25+ n=235 orange. **c)** Single cell RNA-seq (qtSEQ) scatter plot of fold change in gene expression from naïve to TFR (x-axis) and naïve to TFH (y-axis). Colored larger dots indicate p-value <0.05. Unique differentially expressed genes (DEG) for naïve cells are colored blue, TFH green, TFR CD25- purple and shared follicular DEGs are red. **d-f)** Dot plots displaying average gene expression of sorted cells for naïve **(d)**, shared **(e)** and unique **(f)** follicular genes for TFR CD25- and TFH cells. Dot size depicts the percent of cells detecting the gene product and color is average expression.

We had developed a single cell-indexed quantitative and targeted RNA sequencing (qtSEQ) method to evaluate the transcriptional programs of these rare phenotypically complex sub-compartments ^39^. This high-resolution conventional flow cytometry-based sorting method uses 384-well single cell sorting with indexing to track the precise and extended phenotype of the sorted cell through cDNA library formation. In this series of studies, we targeted ∼500 gene products and depending on the cell type sorted, normalized UMI counts per cell ranged from ∼250 to ∼350 molecules detected from our targeted gene set (**Extended Data Fig. 1a-d**). Gene expression was highly specific, when compared to naive B cells with canonical T cell markers *Cd3d, Cd3g, Cd5* at high levels and little to no expression of B cell markers *Cd79a, Cd79b, Cd19* (**Extended Data Fig. 1e-g**). Due to targeted acquisition, single cell qtSEQ provided sensitive representation for the amount and frequency of many genes in these rare lymphocyte sub-compartments.

To define the key molecular programs of activated T_FR_ and T_FH_ cells, we compared each T cell subset to the baseline naive CD4 T cell compartment (**Fig. 1c**) using single cell qtSEQ. Significant changes in the naive T cell program (blue), T_FH_ program (green), CD25^lo^ T_FR_ program (purple) and shared follicular program (red) are displayed. Both T_FH_ and CD25^lo^ T_FR_ cells downregulated genes associated with localization in the T cell zone such as *Ccr7, S1pr1,* and *Sell* as well as transcription factors associated with a quiescent state; *Foxj1, Bcl2,* and *Bach2* (**Fig. 1d**). In addition, T_FH_ and CD25^lo^ T_FR_ cells upregulated a set of shared genes associated with activation and entrance into the follicle including *Cxcr5, Pdcd1, Maf, Icos, Bcl6, Sh2d1a*, and *Slamf6* (**Fig. 1e**). CD25^lo^ T_FR_ cells expressed core regulatory transcripts (*Ctla4, Tnfrsf18, Tnfrsf4, Ikzf2,* and *Havcr2*), ^40^ associated with T_FR_ mediated regulation of IL-1 signaling (*Il1 and Il1r2*) ^37^ and the surface ligand *Sema4c* previously described to direct T_FH_ cells to the GC ^41^(**Fig. 1f**). In comparison to other regulatory subsets CD25^lo^ T_FR_ cells are highest in the expression of follicular markers (**Extended Data Fig. 1h**). In contrast, CD25^hi^ T_FR_ cells express a transcriptional program more related to T_reg_ yet specifically upregulated transcriptional modulators *Irf4* and *Prdm1*. T_reg_ cells expressed markers of localization to the T cell zone (*Ccr7, Sell,* and *Ly6c1*) with naive signatures (*Foxj1* and *Lef1*). However, in this model, we do not detect *Foxp3 mRNA* expression due to the impact of the Thy1.1 reporter on 3’UTR targeting. In contrast, T_FH_ distinctly upregulate molecules associated with co-stimulation and B cell help (*Cd28, Cd44, Il4,* and *Il21*) (**Fig. 1f**) indicating a positive signaling function in the GC. These data emphasize the shared and unique transcriptional components of pre-existing follicular T cell subsets in the spleen at homeostasis before acute PD-1 blockade.

### Acute PD-1 blockade enhances the splenic T_FH_ transcriptional program

Initial studies demonstrated a substantial and selective IgG1 isotype-specific B cell reaction to acute PD-1 blockade ^23^. The response was restricted to the steady-state B cell compartment with no similar effect on antigen-specific induced B cell immunity. CD4 depletion abrogated the impact of treatment, however activated and regulatory CD4 T cell subsets did not change in frequency or total numbers. Hence, we hypothesized that anti PD-1 directly altered the transcriptional programs of GC localized follicular T cell compartments to override type 2 IgG1- specific GC B cell tolerance and produce IgG1 PC and antibodies to gut microbiome.

Using the Foxp3^Thy1.1^ reporter model there were no changes in numbers of activated splenic CD4 T cells and their subsets after acute anti-PD-1 treatment (**Fig. 2a-b**). PD-1 remained blocked with significantly decreased surface expression at day six (**Fig. 2c**) resulting in exaggerated IgG1 PC and IgG1 GC B cell expansion (**Fig. 2d,e**). Even with decreased PD-1 expression, it was possible to sort equivalent compartments of Cxcr5^hi^ T_FH_ cells for single cell qtSEQ from anti PD-1 treated mice (**Extended Data Fig. 2a**). Among multiple enhancements of transcription, we found significant increases in mRNA associated with key T_FH_ attributes such as *Cxcr5* and *Bcl6*, *Sh2d1a* and *Slamf6* with increases in *Il21* and *Cxcr4* (**Fig. 2f**). While PD-1 protein was decreased, mRNA for *Pdcd1* was increased. In addition, anti PD-1 treated T_FH_ cells exhibited significantly higher average UMI counts per cell (**Fig. 2g, left panel**) and upregulated a significant molecular program over steady state T_FH_ (**Fig. 2g, right panel**). Among the broader programmatic changes, we highlight the TCR signaling molecules *Lat* and *Lck,* cell-cell contact proteins *Tnfrsf18, Havcr1,* and *Cd40lg,* as well as the type 2 lineage transcription factor *Gata3* (**Fig. 2h,i**). Further breakdown of significantly upregulated genes by cellular location reveals multiple surface receptors and secreted factors that could augment T_FH_ capacity to provide help to cycling GC B cells (**Extended Data Fig. 2b**). In accordance, gene set enrichment analysis re-iterates significant enhancement of TCR activation pathways, co-stimulation and cytokine signaling (**Extended Data Fig. 2c**). Thus, in the absence of overt T_FH_ cell expansion, there is evidence for enhancement and extension of the pre-existing T_FH_ transcriptional program upon acute PD-1 blockade.

**Figure 2.**
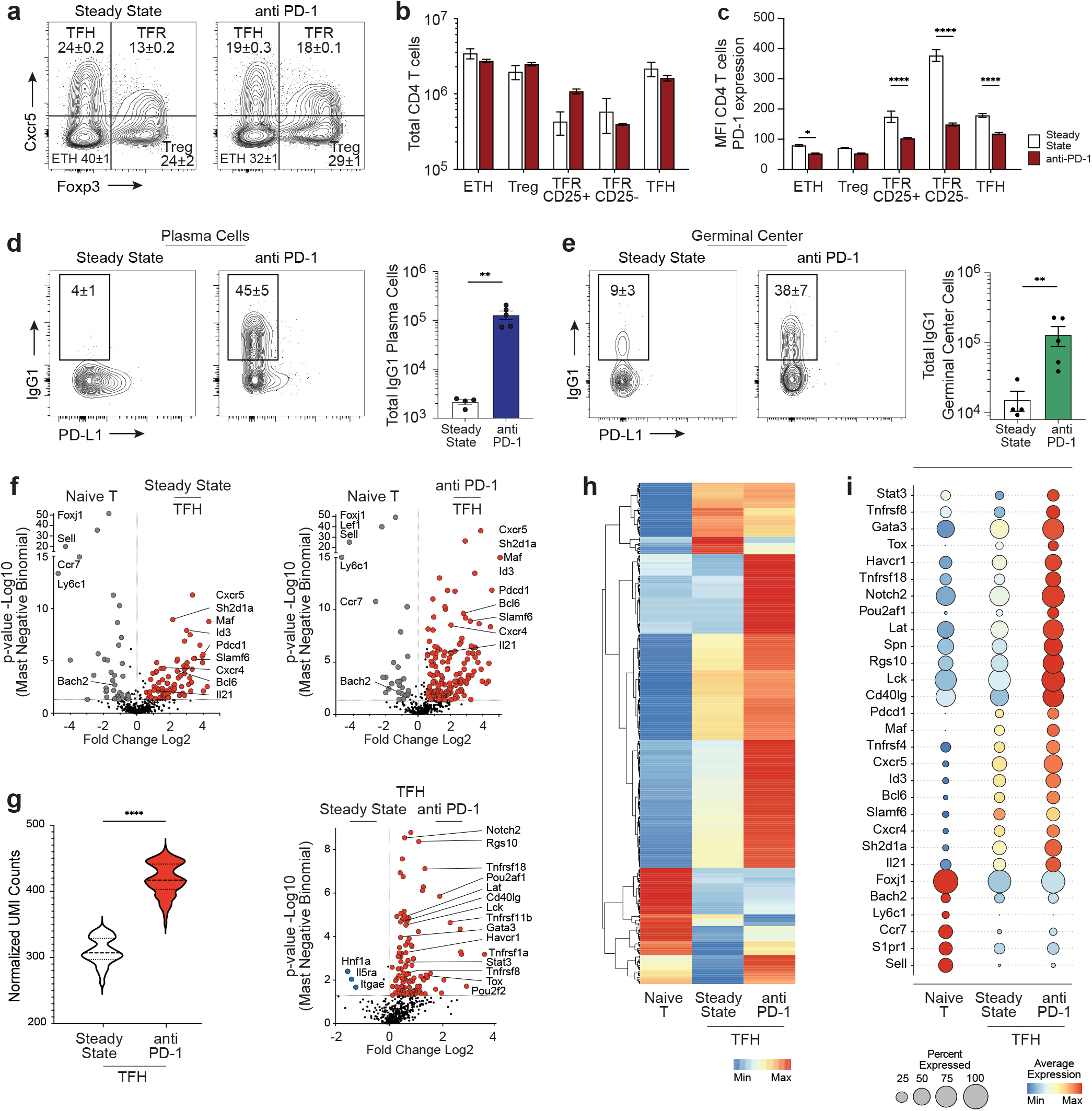
Acute PD-1 blockade enhances the splenic TFH transcriptional program. **a)** Flow cytometry analysis of CD4 T cells (B220-CD8-, CD4+) with intracellular Foxp3 and surface Cxcr5 staining for steady state (n=3) or mice treated 6 days prior with anti PD-1 antibody (clone J43, n=3). **b)** Quantification of total CD4 T cells for each subset and **c)** their PD-1 mean fluorescence intensity (MFI) for steady state (white bars) or anti PD-1 treated (red bars) mice. mean±SEM, p-values *p≤0.05, ****p≤0.0001. **d)** Splenic plasma cells (IgD- IgM- CD138+) for PD- L1 and IgG1 expression with quantification of total IgG1 plasma cell and **e)** germinal center cells (IgD- IgM- CD138- CD19+ CD38- GL7+) with quantification of IgG1 for steady state (n=4) and anti-PD-1 treated mice (n=5). mean±SEM, **p-values <0.01. **f)** Gene expression volcano plots of naïve T cells compared to steady state TFH (left) and naïve T cells to TFH from anti PD-1 treated mice (right). Displayed are Log2 fold change and Mast negative binomial -Log10 p-value. Large and colored dots are gene products that have >1.5 Log2 fold change and/or a p<0.05. **g)** Single cell qtSEQ normalized unique molecular identifier (UMI) counts of TFH cells from steady state mice (n=363) compared to TFH cells from anti PD-1 treated mice (n=418) (left), ****p≤0.0001 and volcano plot of all genes products (right). **h)** Hierarchical clustered averaged single cell heatmap from gene products that had >1.5 Log2 fold change and p<0.05 between naïve T, steady state and anti PD-1 TFH cells with **i)** dot plot of some of the significantly expressed gene products. Color gives average expression and dot size depicts the percent of cells detecting the gene product.

### Checkpoint blockade selectively exaggerates a type 2 GC T_FH_ program

There is growing evidence for functional sub-specialization in the T_FH_ compartment associated with inducing B cell immunity ^30, 36, 42, 43^. To address this issue in the steady-state spleen, uniform manifold approximation and projection (UMAP) dimensionality reduction using gene expression segregates three clusters of T_FH_ (**Fig. 3a,b**). Based on differential gene expression, we designated these subsets as “early T_FH_” (green), “T_FH_1” (purple), and “T_FH_2” (red). Early T_FH_ cells expressed the lowest levels of gene products associated with GC localization such as *Cxcr5, Pdcd1* and *Bcl6* without elements of a type 1 or 2 helper program. In contrast, T_FH_1 cells expressed highest indicators of type 1 function such as *Tbx21* and *Cxcr3* with low expression of the type 2 cytokine *Il4*. T_FH_2 cells expressed the highest levels of type 2 gene products such as *Maf, Gata3, Notch1* and *Il1rl1* and were also highest in the relative expression of genes affiliated with localization in the GC (*Pdcd1, Cxcr5,* and *Il21*). These designations were supported by corresponding UMAP clustering based on key phenotypic markers that also revealed three separable T_FH_ cell clusters (**Extended Data Fig. 3a**). Most notably, a T_FH_1 subset (purple) expressed highest levels of CXCR3/*Cxcr3* protein/message and clearly separates from a T_FH_2 cluster (red) expressing highest levels of PD1 and CXCR5 that was defined at the transcriptional level by those markers as well as *Il4, Maf,* and *Gata3* (**Extended Data Fig. 3b**).

**Figure 3.**
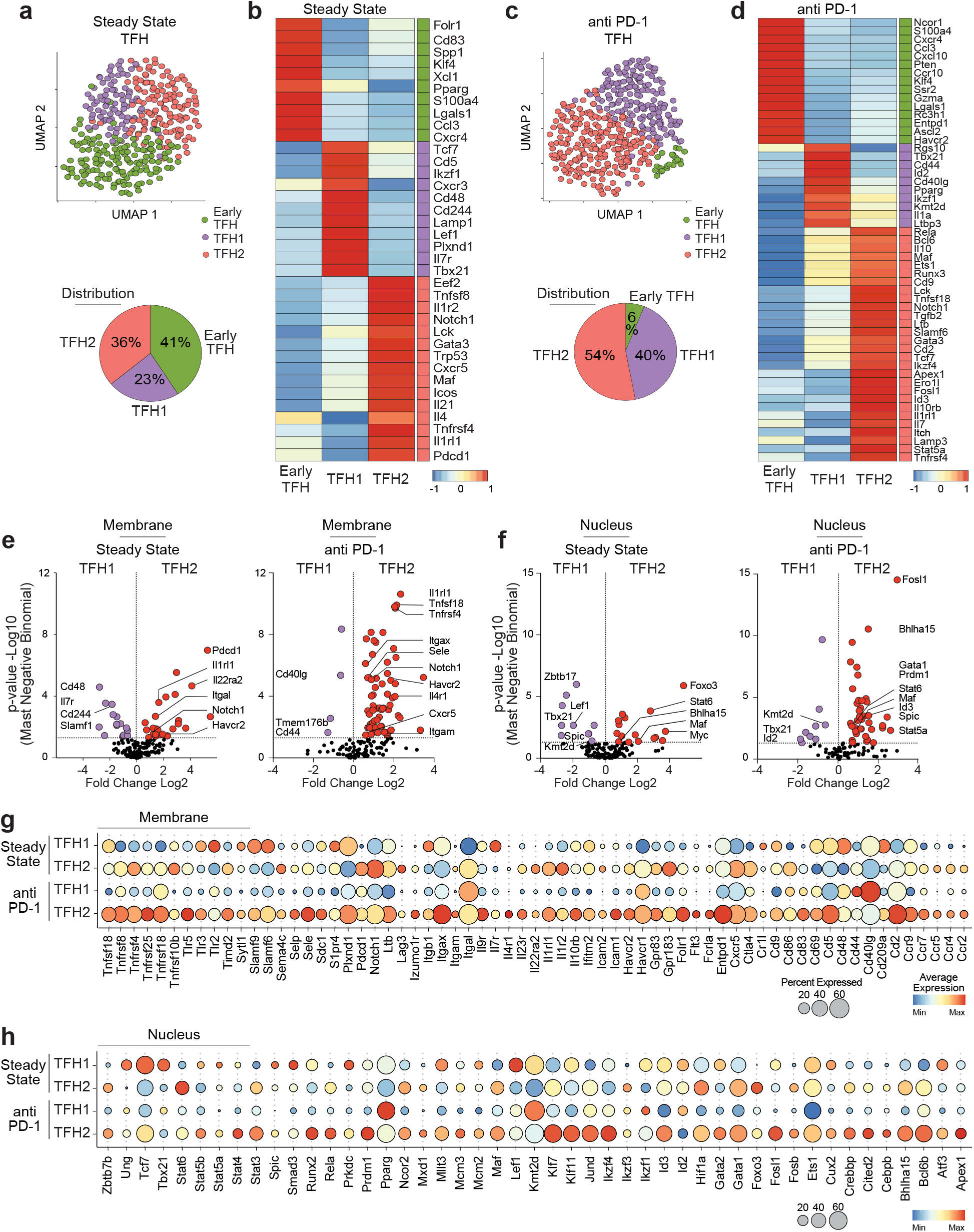
Checkpoint blockade selectively exaggerates a Type 2 GC TFH program. **a)** Steady state TFH cells unbiased Uniform Manifold Approximation and Projection (UMAP) cell clusters (n=363, early TFH colored green, TFH1 purple, TFH2 salmon) with a distribution pie chart and **b)** single cell averaged heatmap of the top significant gene products (>1.5 Log2 fold change and p<0.05) differentiating each cluster (using Seurat FindMarkers function). **c)** UMAP clusters of anti PD-1 treated TFH cells (n=418), distribution pie chart and **d)** averaged single cell heatmap of the top differentiating gene products for each cluster. **e)** Volcano plots comparing genes encoding proteins localized in the cellular membrane or **f)** the nucleus from the TFH1 cluster (n=84) and TFH2 cluster (n=129) at steady state or in the same clusters after PD-1 blockade (anti PD-1: TFH1 cluster n=167, TFH2 n=228). Large and colored dots were considered significant (>1.5 Log2 fold change and p<0.05). **g)** Dot plot of all significantly expressed gene products localized in the membrane and **h)** nucleus for TFH1 and TFH 2 cells from steady state or anti PD-1 treated mice. Color shows average expression and dot size depicts the percent of cells detecting the gene product.

Based on similar patterns of differential gene expression, we can readily designate fractions of “T_FH_1” (purple), and “T_FH_2” (red) clusters within the T_FH_ compartment after acute PD- 1 blockade with a substantial decrease in the “early T_FH_” subset (green) (**Fig. 3c,d**). While the resultant fractions of T_FH_1 and T_FH_2 were similar in number, the transcriptional program of the T_FH_2 was selectively exaggerated after treatment (**Extended Data Fig. 3c,d**). While many of the steady-state elements of this program (*Cxcr5, Slamf6, Bcl6, Gata3, Maf, Il4, Il21*) were further enhanced with treatment (**Fig. 3d**), there were extended transcriptional changes selectively exaggerated in this T_FH_2 subset (**Fig. 3e-h; Extended Data Fig. 3e-g**). T_FH_2 cells upregulated multiple members of the TNF superfamily of receptors and ligands (*Tnfsf8, Tnfsf18, Tnfsf13, Tnfrsf8,* and *Tnfrsf4)* with many examples of increased cell-cell adhesion (*Cd44, Sdc1, Icam1, Icam2*, *Sele, Itgax,* and *Itgal*), modifiers of cell interactions (*Plxd1, Sem4c*, *Havcr1, Havcr2, Timd2*), metabolism (*Entpd1, Gpr83, Gpr183, Folr1*) increased cytokine receptors (*Il4r, Il7r, Il9r, Il10rb*), costimulatory molecules (*Cd28, Cd2, Cd40lg, Cd69*) and other surface interactors capable of altering cell function (Ctla4, Lag3, Notch1). Hence, transcriptional increases associated with molecules expressed on the cell membrane (**Fig. 3e,g**) were abundantly released after PD-1 blockade and selectively exaggerated in the T_FH_2 compartment. PD-1 blockade also selectively enhanced major transcriptional regulators in the T_FH_2 compartment in comparison to T_FH_1 cells (**Fig. 3f,h**). There are increases in multiple Stat family members (*Stat3, Stat4, Stat5a, Stat5b*), *Gata1* and *Gata2* with a switch from *Id2* to *Id3* prevalence, and increases in multiple other in major transcription factors known to modify lymphocyte function (*Runx2, Klf7, Klf11, Ikzf4, Prdm1, Bhlha15, Bcl6b*). Changes in expression for secreted molecules (*Tgfb1, Il7, Ebi3, Tnfs13, Il6, Tslp, Tnsf8*) and those residing in the cytosol (*Lamp1, Eef2, S100a4, Lck, Atf6*) (**Extended Data Fig. 3e-g**) further exemplify the selective targeting of the T_FH_2 transcriptional program by acute PD-1 blockade.

To consider how these molecules interact to enhance immune function, Metascape ^44^ was used to visualize shared and unique gene expression patterns across T_FH_1 and T_FH_2 (**Fig. 4a**). After PD-1 blockade, the exaggerated T_FH_2 unique program increases at the expense of genes expressed in the T_FH_1 compartment. Network density plots and their elaboration in pathway displays using MCODE ^45^ highlight the most connected protein networks and their associated gene ontology (GO) pathways (**Fig. 4b,c**). While network density is low before treatment in the T_FH_1 compartment, this increases with treatment to significantly influence lymphocyte differentiation and chemotaxis (**Fig. 4b**). However, the T_FH_2 network accessed here is already dense before treatment with evidence of PD1 signaling and elements of T_FH_2 pathway activation. Upon release of the PD-1 checkpoint, the plot displays significant exaggeration and elaboration of pathways involving T cell activation, cytokine signaling and cellular differentiation (**Fig. 4c**). Hence, the network analysis re-iterates the selective targeting of a pre-existing T_FH_2 transcriptional program that is enhanced and exaggerated upon the release of a steady-state PD- 1 GC checkpoint.

**Figure 4.**
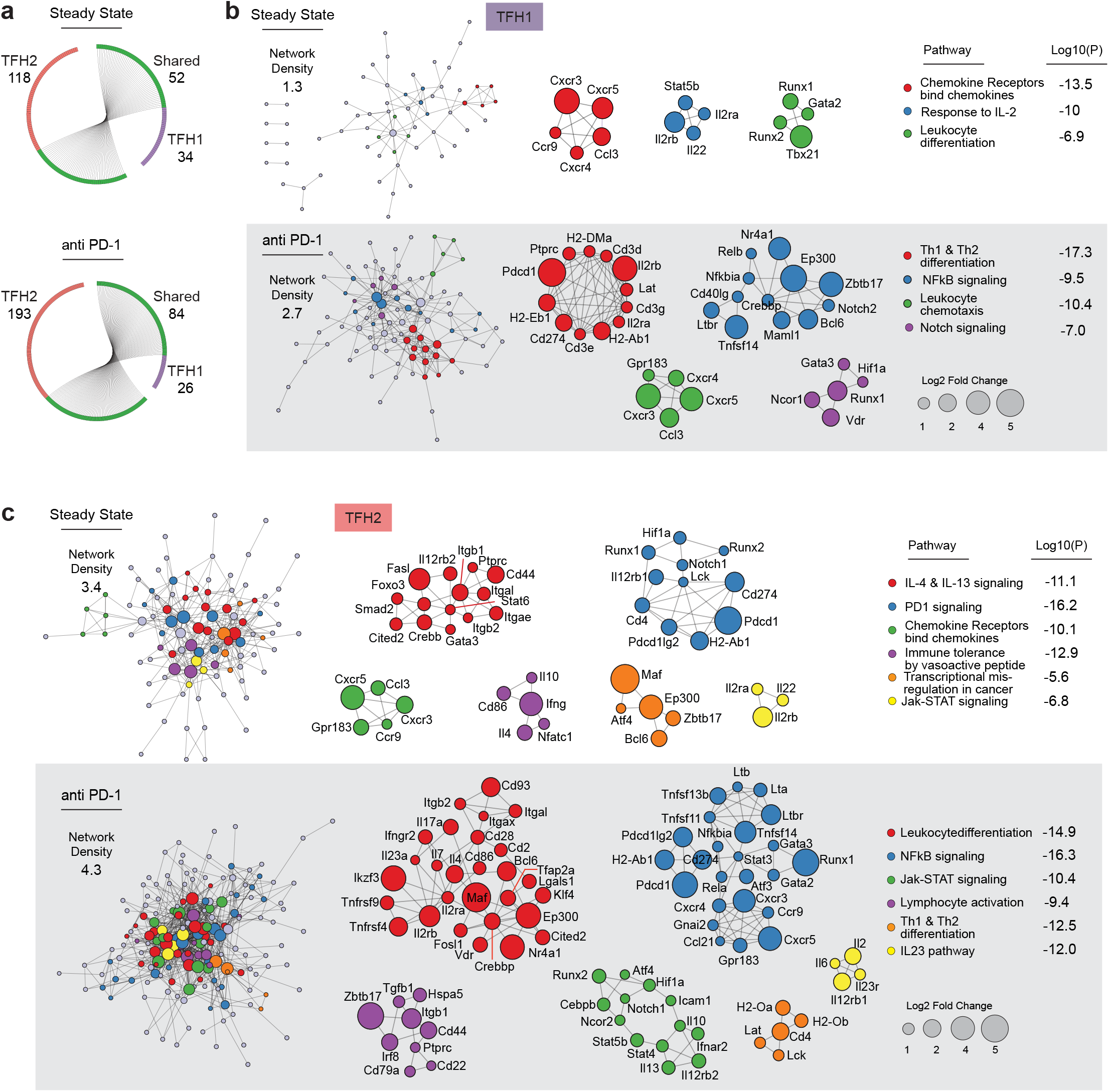
Predicted protein interactome network of the enhanced Type 2 GC TFH program. **a)** Circos visualization of TFH1 or TFH2 cells compared to naïve T at steady state (upper) or following PD-1 blockade (lower) significantly differentially expressed gene products (>1.5 Log2 fold change and p<0.05). Upregulation of shared (green edge with connected black lines) and unique (no line) gene products for TFH1 (purple edge) or TFH2 (salmon edge) cells. Number of gene products is indicated for shared, TFH1 and TFH2. **b, c)** Metascape visualization of the interactome network form by **b)** TFH1 **c)** TFH2 steady state genes (upper panel) and after PD-1 blockade (lower grey panel). Network density is the number of edges divided by the number of nodes. The size of each node positively correlates with number of edges it connects. Cluster complexes are colored according to their functional identities or enriched gene ontology pathway. The molecular complex detection (MCODE) algorithm identifies densely connected protein networks that are associated with a broader functional unit (min size = 3 nodes), are displayed with mouse gene names and drawn from colored dots in the network. Dot size reflects the Log2 fold change upregulated from naïve to TFH1 or TFH2. Log10 p-values are displayed.

### Divergent transcriptional program of GC-localized T_FR_ cells

Checkpoint inhibitors can exert control of effector function through regulatory T cell subsets, with for example T_reg_ specific PD1 deficiency resulting in enhanced suppressive capacity^46^. UMAP display of sorted Foxp3 expressing regulatory T cell compartments from the steady-state spleen segregates 3 subsets of T_FR_ from conventional Treg cells (**Fig. 5a**). As discussed earlier, we can designate an ’early’ CD25^hi^ T_FR_ subset (green) that expresses lowest levels of Cxcr5 and PD-1 and likely to be localized in the follicular regions, but outside the GC reaction. There was also a small but distinct Cxcr3 expressing T_FR_ compartment (purple) that expressed intermediate levels of Cxcr5 and PD-1. In contrast, the majority of the remaining T_FR_ compartment expressed highest level of Cxcr5 and PD-1 (red) and likely to be localized within the GC. We refer to this subset as ’GC-T_FR_’ and scrutinize this subset more closely as potential suppressors of GC B and T_FH_ function at homeostasis. We could readily distinguish all four subtypes of regulatory T cells after PD-1 blockade, even with decreased PD-1 levels (**Extended Data Fig. 4a**). Comparative gene expression analysis helped to reinforce the divergent transcriptional programs expressed by these phenotypically separable regulatory compartments (**Fig. 5b**). The Cxcr3^+^ T_FR_ cells also expressed highest levels of Cxcr3 and Lgals1 gene expression. Overall, the divergent gene expression programs of all three T_FR_ cell fractions displayed minor changes after PD-1 blockade (**Extended Data Fig. 4b-f; Fig. 5c-f**).

**Figure 5.**
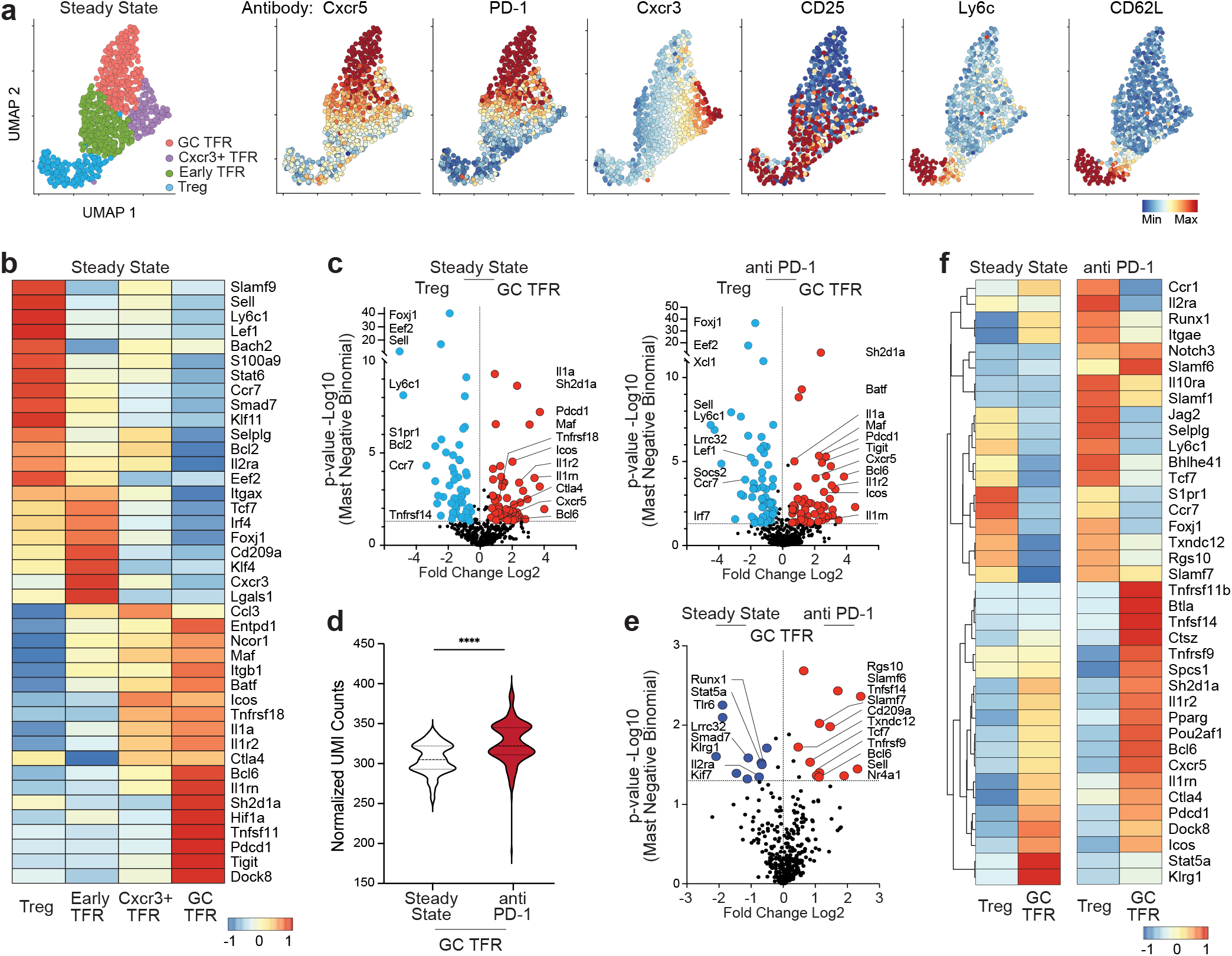
Impact of PD-1 blockade on TFR cells. **a)** Foxp3^Thy1.1^ reporter mice UMAP flow cytometry analysis of steady state sorted (see Fig 1b) CD4 T (CD4+ TCRb+), regulatory Treg (Thy1.1+ Cxcr5-), TFR CD25- (Thy1.1+, Cxcr5+, CD25-) and TFR CD25+ (Thy1.1+, Cxcr5+, CD25+) cells. Five parameters (Cxcr5, PD-1, Cxcr3, Ly6c, CD62L) were used to cluster the cells (left). The 4 Foxp3 (Thy1.1+) clusters identified Treg (colored blue), early TFR (green), Cxcr3+ TFR (purple) and GC TFR (salmon) cells. Distribution of individual antibodies Cxcr5, PD-1, Cxcr3, CD25, Ly6c, CD62L are overlaid with each fluorophore (right panels) intensity from minimum (blue) to maximum (red). **b)** Single cell qtSEQ averaged heatmap of the top significant marker genes differentiating each regulatory cell cluster from steady state mice (n=663). **c)** Gene expression volcano plots of steady state Treg (n=133) compared to GC TFR cells (n=221) (left) and anti PD-1 Treg (n=117) to GC TFR cells (n=272) (right). Displayed are Log2 fold change and Mast negative binomial -Log10 p-value. Large and colored dots are gene products that have >1.5 Log2 fold change and/or a p<0.05. **d)** Normalized unique molecular identifier (UMI) counts of GC TFR cells from steady state to anti PD-1 treated mice (****p≤0.0001) and **e)** volcano plot of the gene products from GC TFR cells steady state and anti PD-1 treated mice. **f)** Hierarchical clustered averaged single cell heatmap from gene products that had >1.5 Log2 fold change and p<0.05 between Treg and GC TFR cells from steady state and anti PD-1 mice.

While changes were small, it was of interest to consider the impact of PD-1 blockade within the GC T_FR_ compartment predicted to exert more local constraint on type 2 IgG1 GC B cells at homeostasis. While steady state GC T_FR_ cells were notably distinct from steady state T_reg_ cells, most elements of this divergent molecular program were not enhanced after PD1 blockade (**Fig. 5c**). Averaged UMI counts were significantly increased but only slightly higher after treatment (**Fig. 5d**). The composition of this program did vary after treatment with GC T_FR_ significantly upregulating follicular genes such as *Bcl6* and *Tcf7* along with cell surface effector molecules *Tnfrsf9, Tnfsf14,* and *Slamf6* (**Fig. 5e,f**). These alterations potentially impact suppressive function in the GC but also downregulating potential regulatory effector molecules such as *Lrrc32* and Klrg1 makes interpretation difficult. In conclusion, even the GC T_FR_ cells were only moderately impacted by PD-1 blockade compared to the enhanced transcriptional programs seen in T_FH_2 cells.

### PD-1 blockade alters the regulatory balance of GC-T_FR_ and T_FH_2 subsets

The extent to which T_FR_ cells directly interact with GC B cells or T_FH_ cells *in vivo* remains unclear, although *in vitro* co-cultures have shown the T_FR_ cells are able to exert suppressive control over both subsets ^47, 48^. It is likely the balance of effector molecule expression on either helper or regulatory follicular cells would play a role in shaping GC B cell fate and function. To investigate how PD-1 blockade might alter the regulatory potential between GC T_FR_ and T_FH_2 cells we compared expression of key effector molecules between the two subsets.

GC T_FR_ cells at steady state expressed a regulatory signature over T_FH_2 cells (*Ctla4, Cd72, Tnfrsf9, Sema4c, Il1r2, Il12rb1,* and *Itgae*) that was maintained after PD-1 blockade (**Fig. 6a,b**). However, GC T_FR_ cells lost expression of *Klrg1* and *Lilrb4* while T_FH_2 cells upregulated multiple transcripts encoding cell-cell contact proteins previously exclusively expressed by GC T_FR_ cells at steady state (*Cd83, Tnfrsf18, Cd244,* and *Cd48*). The T_FH_2 subset also encoded a helper signature of membrane proteins that was enhanced after anti-PD1 blockade (*Notch1, Gpr183, Cd27, Cd44,* and *Itgam*). Significantly PD-1 blockade induced T_FH_2 cells to upregulate numerous membrane proteins that were not expressed over GC T_FR_ cells at steady state including multiple soluble factor receptors (*Il10rb, Il22ra1,* and *Il9r*) as well as several TNF family members (*Tnfsf11b, Tnfrsf13c, Tnfrsf14, Tnfrsf17,* and *Tnfsf18*). PD-1 blockade also altered the balance of secreted effector molecules between follicular helper and regulatory subsets (**Fig. 6c,d**). GC T_FR_ cells maintained significant expression of *Ccl5* and the regulatory cytokine *Fgl2* which has been linked with T_reg_ mediated suppression of Th2 cytokine production ^49^. In contrast, T_FH_2 cells maintained a helper signature of secreted proteins (*Il21, Il15, Tnfsf8, Tnfsf13* and *Tgfb3*) and upregulated a *de novo* secreted signature (*Tgfb3, Il3, Spp1, Tgfb1, Il17a* and *Xcl1*) after anti PD- 1 blockade. In similar fashion T_FH_2 cells exhibited a strongly enhanced program of transcriptional regulators (**Extended Data Fig. 5a,b**) and intracellular signaling molecules **(Extended Data Fig. 5c)** over GC T_FR_ cells following PD-1 blockade. Taken together, these data indicate that while PD- 1 blockade does impact GC T_FR_ cells, it is the T_FH_2 program that is significantly augmented at the expense of the regulatory subset releasing positive effector function and driving maturation of IgG1 B cells within the steady-state GC microenvironment.

**Figure 6.**
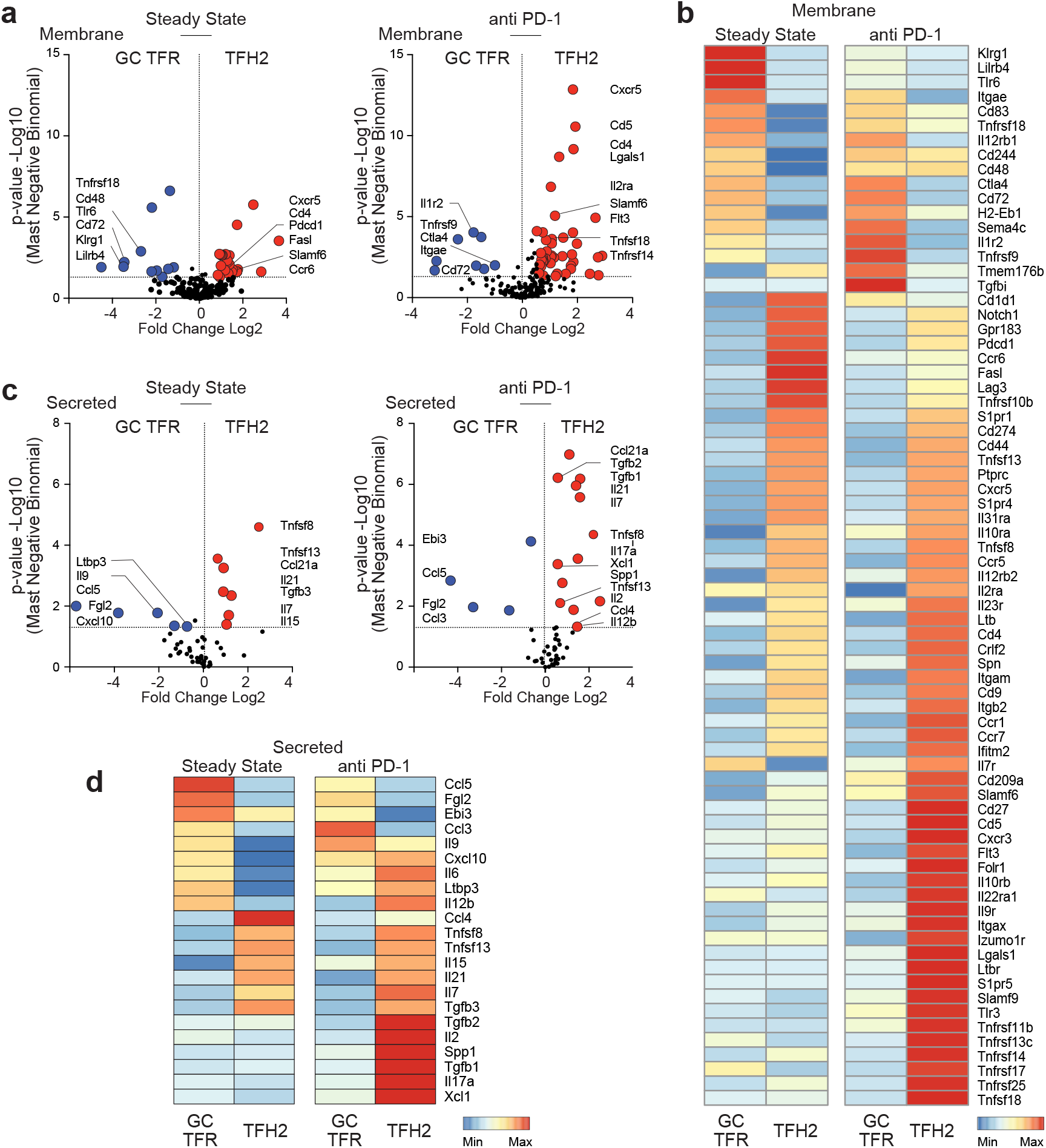
PD-1 blockade shifts the GC regulatory balance. **a)** Single cell gene expression volcano plots comparing genes encoding proteins localized in the cellular membrane from steady state GC TFR (n=221) and TFH2 cells (n=129) (left) and after PD- 1 blockade (GC TFR n=272, TFH2 n=228 cells) (right). Large and colored dots were considered significant (>1.5 Log2 fold change and p<0.05). **b)** Averaged single cell heatmap of the significant cell membrane products identified in a). **c)** Volcano plots comparing steady state and anti PD-1 treated GC TFR and TFH2 cells from secreted gene products with **d)** averaged single cell heatmap of the significant gene products identified in c).

### Enhanced T_FH_2 to IgG1 GC B inter-cellular contact underpins GC maturation

To assess the variety of inter-cellular contacts involved in regulating the type 2 GC B cell compartment, we compared changes in the GC T_FR_ and T_FH_2 transcriptional programs to those from IgG1 GC B cells ^23^. Connectome ^50^ is used to visualize predicted ligand receptor interactions between all three cell types before and after PD-1 blockade. At steady state T_FH_2 cells had limited potential to influence GC IgG1 B cells with capacity to interact through *Tnfrsf13-Tnfrsf13b* and Icam1-integrin complexes *Itgax* and *Itgal* (**Fig. 7a left panel, Extended Data Fig. 6a**). Similarly, GC T_FR_ cells could contact steady-state IgG1 GC B cells using *Tnfrsf18-Tnfsf18.* However, after PD1 blockade the number of predicted interactions between T_FH_2 and GC IgG1 B cells increased substantially compared to those with GC T_FR_ cells (**Fig. 7a right panel, Extended Data Fig. 6b**). Enhanced cytokines (*Il2, Il17, Il23a*) with corresponding receptor sub-units (*Il2rb, Il17ra, Il23r*) and chemokines (*Ccl3, Ccl19*) and their receptors (*Ccr1, Ccr4, Cxcr3*) establish new means of driving T-B interactions that may enhance IgG1 GC B cell activation. Changes in T_FH_2-IgG1 GC B cell connections between *Notch1/2* and *Jag1/2* could enhance signaling pathways thought to drive T_FH_ differentiation, T cell activation, and *Il4/Gata3* production ^51^.

**Figure 7.**
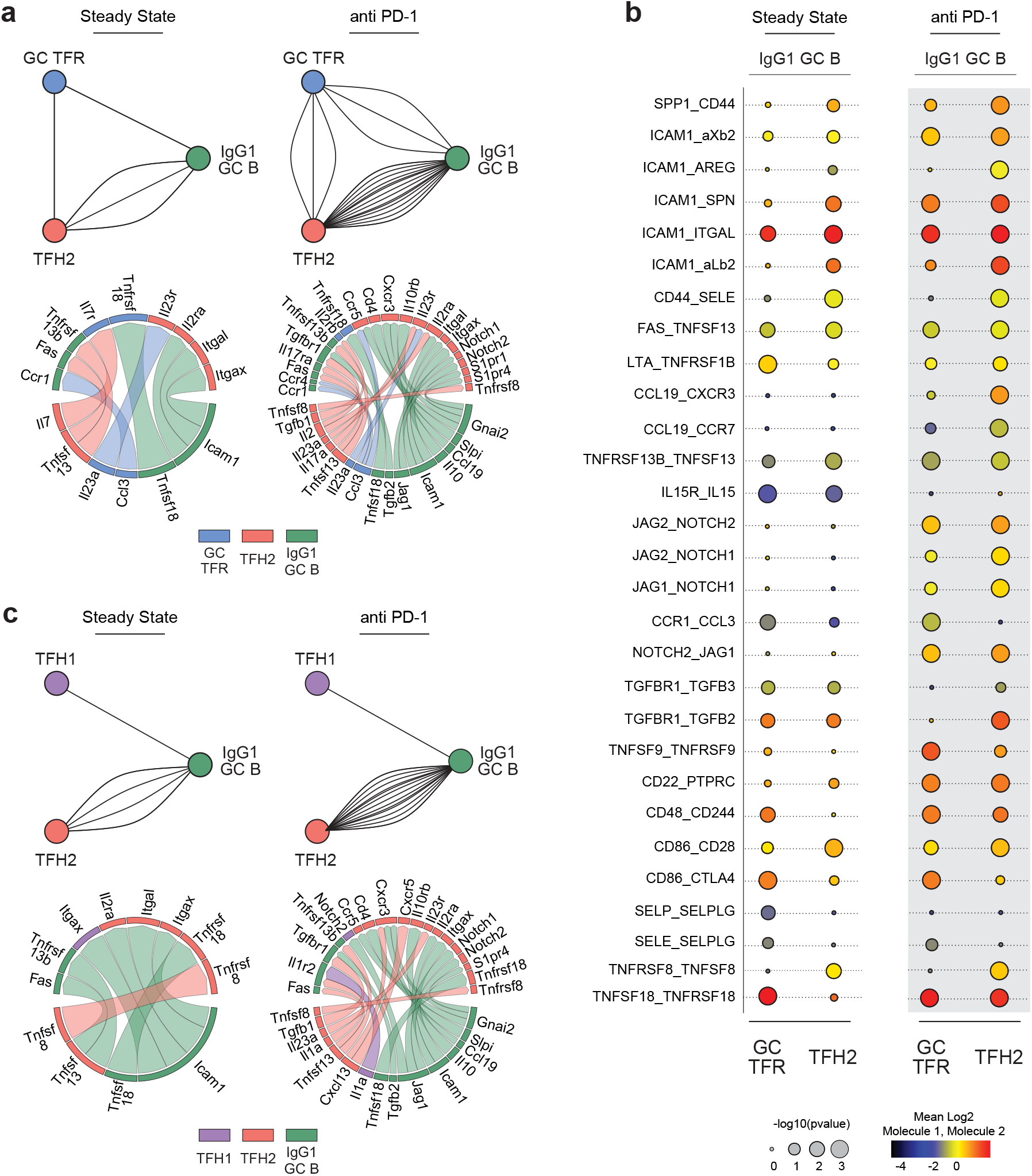
Enhanced T_FH_2 to IgG1 GC B inter-cellular contact underpins GC maturation. **a)** Single cell Connectome cell-cell connectivity patterns by ligand-receptor mapping of GC TFR (blue), TFH2 (salmon) and IgG1 GC B (green) cells from steady state (left) and anti PD-1 treated mice (right). Each line represents a predicted ligand-receptor interaction (upper) with details of each individual ligand-receptor interaction (below). Connections are filtered by significance (p<0.05) and minimum fraction of cells expressing a gene product (>10%). **b)** Cell-cell communication facilitated by receptor-ligand contacts between IgG1 GC B cells and either GC TFR or TFH2 cells from steady state and PD-1 blockade using CellPhoneDB. Receptor ligand interactions are listed from left (predominately expressed by B cells) to right (predominately expressed by T cells). P-values are indicated by dot size, and color indicates mean Log2 average expression level of interacting molecules. **c)** Connectome cell-cell network by ligand-receptor mapping of TFH1 (purple), TFH2 (salmon) and IgG1 GC B (green) cells from steady state (left) and anti PD-1 treated mice (right) as described in a).

Complementary receptor-ligand analysis using CellphoneDB ^52^ re-iterates the broad observations above, indicating PD-1 blockade promoting the highest numbers of intercellular connections T_FH_2-GC IgG1 B contacts (**Extended Data Fig. 6c**). This analysis highlighted additional interactions beyond *Tnfrsf18-Tnfsf18*, including *Tnfsf8/Tnfrsf8* and *Fas-Tnfsf13* (**Fig. 7b**). Some molecules upregulated in the T_FH_2 compartment such as SLAM family member *Cd48* and *Cd244* were exclusive to GC T_FR_ before treatment. Multiple adhesion/integrin interactions were increased through upregulation of *Icam1* on IgG1 GC B cells and increases in multiple counter-receptors on T_FH_2 cells (including *Itgal*, *Spn* and *Areg)*, while GC T_FR_ cells lost potential B cell contacts: *Selp-Selplg, Il15-Il15ra,* and TGF beta signaling. Using the same analysis, we directly compared predicted T_FH_2 and T_FH_1 connections to GC IgG1 B cells and re-emphasize the selective increase in T_FH_2 contacts after PD-1 blockade (**Fig. 7c**). Hence, combining these T-B datasets to focus on predicted inter-cellular contacts, we identify multiple pathways that are enhanced by PD-1 blockade with potential to activate the GC B cell cycle and release IgG1 isotype-specific adaptive B cell tolerance.

## DISCUSSION

To establish and maintain homeostasis, adaptive immunity must enforce tolerance against environmental antigens. We recently demonstrated a PD-1 dependent immune checkpoint within steady-state splenic GC that constrains the maturation of type 2 IgG1 isotype-specific GC B cells^23^. The IgG1 selective block in splenic GC B cell cycling further restricts the production of these IgG1 antibody specificities at homeostasis. Upon PD-1 blockade, released specificities were reactive to gut microbial antigens with no evidence for autoimmunity or binding to dietary antigens in this model. Extended studies on induced immunity to foreign antigen revealed no such impact, strongly indicating that this PD-1 checkpoint selectively regulates adaptive B cell tolerance and not immunity.

Here, we define the transcriptional program of multiple PD-1 expressing follicular T cell compartments to identify the direct CD4 T cell target of acute PD-1 blockade. Using single cell-indexed RNA sequencing, we find pre-existing subsets of splenic T_FH_ and T_FR_ compartments and identify type 2 T_FH_ cells as the central target of PD-1 blockade. An exaggerated T_FH_2 transcriptional program overcomes local GC T_FR_ regulatory processes to drive TCR activation, promote cytokine signaling with enhanced cell fate modifying surface contacts with IgG1 GC B cells. Hence, acute PD-1 blockade selectively induces T_FH_2 transcriptional re-programming to release type 2 IgG1 GC adaptive B cell tolerance and preferentially promote IgG1 PC formation and enhanced antibody production.

The adaptive immune system emerges and persists in layers of complementary and opposing functions. Here, we consider the follicular CD4 T cell compartments that sub-specialize to induce and regulate adaptive B cell immunity ^5, 6^. Inducer T_FH_ cells bifurcate from conventional ETH cells based on the antigen-presenting inflammatory context of initial antigen experience from naive antigen-specific precursor T_H_ cells. Effector T_FH_ cells can then influence multiple stages of activated antigen-specific B cell fate, including the earliest PC versus GC entry decision ^53,54,55,56,57^. In contrast, T_FR_ cells are recruited from pre-existing Treg compartments that were produced through selection on self-peptide MHCII complexes during thymic development ^7, 8^. Therefore, the T_FR_ compartment is self-reactive and influences B cell fate in non-cognate ways through secreted and cell contact associated actions. In the current study, we focus on the steady-state splenic environment that contains PC, GC B cells and memory B cells of all antibody isotypes from IgM to all downstream IgG subclasses ^23^. From these initial studies, we know that some of the steady-state GC B cells, IgG PC compartments and circulating antibodies express anti-microbiome specificities. Therefore, pre-existing splenic follicular T cell compartments at homeostasis must support, not only previous foreign immune challenge, but also deploy adaptive B cell tolerance mechanisms.

Inducing antibody class switch in activated B cells is an integral component of early T_FH_ function that is thought to largely precede GC entry ^53^. In this context, T_FH_ function can be segregated in principle across the major immune response modules; type 1 inflammatory IgG2 subclass, type 2 IgG1 anti-inflammatory and type 3 IgA mucosal antibody ^28, 29, 31, 32^. We and others have proposed this broad organization for T_FH_ induction ^36, 58, 59^ that may extend to T_FR_ regulation ^54, 60^. We have demonstrated a separable requirement for T-bet in the survival and reactivation of type 1 IgG2a expressing memory B cells that implies existence of a memory T_FH_1 isotype-specific inducer ^33^. We have recently extended these studies to demonstrate the differential isotype-specific programming of PC compartments, both antigen-specific and pre-existing in the steady-state ^31^. Furthermore, we provide evidence for the segregation of T_FH_1 and T_FH_2 compartments based on divergent transcriptional programs deployed over time to modify B cell antibody isotype^30^. Recent discovery of T_FH_13 cells that separably regulate IgE responses ^36^ and evidence for separable requirements for IgA immunity ^58^ further supports this conceptual framework. In this context, the T_FH_1 versus T_FH_2 designation of pre-existing splenic subsets in the current study is readily reconcilable, while the ’early’ T_FH_ assignation remains poorly resolved.

T_FH_ induction of memory B cell responses at recall may also be segregated to activate, expand and differentiate already switched memory B cells through a memory response PC versus GC fate decision ^61^. Therefore, at steady-state pre-existing T_FH_ compartments either support ongoing effector function, exist as persistent and quiescent memory T_FH_ compartments or display a tolerance inducing program to constrain B cell immunity ^23^. In the initial study, we demonstrated that the pre-existing type 2 IgG1 GC B cell compartment, but not the type 1 IgG2 GC B cell compartment, was heavily restricted to the light zone of the GC reaction. Thus, we predicted at least some component of the pre-existing splenic T_FH_ compartment would display this tolerance inducing behavior. Hence, the transcriptional program in the T_FH_2 compartment was of particular interest. Multiple molecular pathways enhanced by treatment can be considered in the context of known PD-1 action dysregulating constraints on TCR activation through enhanced co-stimulation, cytokine signaling, cell-cell adhesion ^62, 63^. These changes alone provide multiple means for enhancing contact with IgG1 GC B cells to drive dark zone entry and the ongoing GC cycle for the type 2 IgG1 isotype.

The selective targeting of IgG1 remains the central observation and provides a compelling example of CD4 driven isotype-specific regulation of the antigen-experienced B cell compartment. In the previous study, we proposed that the IgG1 pathway acts as a reservoir for IgE ^64, 65^. Therefore, restricting affinity maturation of IgG1 antibody selectively constrained the specificities that may progress into the IgE compartment. Here, we provide evidence that selective PD-1 targeting occurs more directly at the level of the pre-existing T_FH_2 compartment. This is further reconciled by the higher level of PD-1 expressed by the T_FH_2 subset in the steady-state, providing a simple means for differential impact of blockade. However, local and focal changes in the distribution of the PD-1 ligands have not been studied in the context of a pre-existing or developing GC reaction and may provide important contextual information.

Recent studies begin to uncover specific mechanisms that drive affinity maturation ^66,67,68^. However, functional interactions that enable productive T_FH_-GC B cell cognate contact and the rules that control post-GC cell fate remain complex and difficult to access experimentally. It is also plausible that all GC are not the same. We provide evidence that antibody isotype can distribute unevenly in a developing primary response GC as one indication of variability ^30^. It is also possible that GC containing long-term antigen depots, or readily renewable depots of microbial antigens, are fundamentally different to de novo developing and short-term resolving GC ^69^. In this scenario, both the stromal microenvironment and the resident T_FH_ populations may be primed and programmed in qualitatively or quantitatively distinct ways that segregate the molecular response to PD-1 and its subsequent release at blockade.

The subset separation of steady-state regulatory T cells is also consistent with current thinking. We identify molecules likely involved in differentiation from T_reg_ to CD25^hi^ T_FR_ to a fully matured CD25^lo^ T_FR_ and confirm multiple components T_FR_ transcriptional programs identified using bulk RNA-SEQ ^11, 37, 70, 71^. Here, among the CD25^lo^ compartment, small numbers of Cxcr3^+^ T_FR_ were described with lower propensity for GC localization. We pursued focused analysis of the Cxcr5^hi^ PD1^hi^ Bcl6^hi^ expressing compartment most reasonably designated GC T_FR_. Evidence of direct *in vivo* T_FR_ cell contact with GC B cells is lacking and cognate contact may not frequently occur in the context of foreign antigen ^7, 72^. However, T_FR_ cells have been shown to employ non-specific mechanisms of suppression to regulate GCs. The secretion of neuritin ^72^, expression of CTLA-4 ^73, 74^ and control of IL-1 signaling ^37^ have all been associated with T_FR_ regulation of GCs.

Nevertheless, under PD-1 blockade that leads to overt and substantial type IgG1 GC B cell dysregulation, there was very little change in the GC T_FR_ program.

The enhancement and exaggeration of a T_FH_2 transcriptional program with little change in local attributes of GC T_FR_ combine to dysregulate the homeostatic type 2 GC B cell balance. All compartments are predicted to exist in humans and can be susceptible to PD-1 checkpoint blockade used in the clinic. Anti-inflammatory murine IgG1 is most similar in function to IgG4 in humans ^35^. Recent studies connect T_FH_ and B cell activation as a strong predictor of checkpoint inhibitor efficacy ^75^. Similarly, microbiome composition is linked to checkpoint inhibitor efficacy ^76,77,78^. The impact of checkpoint blockade will depend on the composition of antibody specificities that exist in the individual’s GC reaction at steady state. For example, autoimmune susceptible individuals may elaborate autoantibody responses upon release of tolerant specificities. Allergy prone individuals may similarly exaggerate an IgG4 response that accelerates IgE antibody production. It remains intriguing to examine how these mechanisms of T follicular cell control of germinal center dynamics and insight into the emergent properties of isotype-specific immune regulation operate in human health and disease.

## MATERIALS and METHODS

### Mice

Foxp3-IRES^Thy1.1^ Reporter mice ^79^ obtained from Ye Zheng (Salk Institute for Biological Studies) were bred and housed under specific pathogen-free (SPF) conditions at The Scripps Research Institute. Experimental protocols were approved by Institutional Animal Care and Use Committee at The Scripps Research Institute. Male 8-20 week old mice were euthanized with isoflurane prior to sample collection and analysis.

### In vivo antibody treatments

300μg anti-mouse PD-1 antibody (clone J43, BioXcell) was administered intraperitoneal (IP) for in vivo blockade of PD-1 and mice were analyzed 6 days post treatment. Steady state mice received no treatment.

### Flow cytometry

As previously described ^61^, single-cell suspensions were made from the spleen. Processing of cells and flow cytometry staining were performed on ice. Briefly, cells were lysed (ACK red cell lysis buffer, Gibco), passed through a 70μm cell strainer (Fisher Scientific), washed, and resuspended in ice cold phenol red-free DMEM (Gibco) with 2% (vol/vol) Fetal Calf Serum FACS buffer at 4 x 10^8^ cells/mL.

For surface staining, cells were incubated for 5 mins with 2% (vol/vol) normal mouse and normal rat serum (Jackson ImmunoResearch Laboratories), prior to staining with a 2X master mix containing fluorophore-labeled antibodies at pre-titrated concentrations in Brilliant Violet Staining Buffer (BD Biosciences), and added to suspended cells at a final staining concentration of 2 x 10^8^ cells/mL for 30-45 min. Cells were washed and resuspended in FACS buffer before flow cytometry analysis and sorting. For surface staining of IgG1, non-specific binding was first blocked with CD16/32 (clone 2.4G2, BioXcell) for 5 mins prior to adding the master mix of fluorophore-labeled antibodies.

For Foxp3 intracellular staining, surface staining was completed first. Cells were incubated with Invitrogen, eBioscience Fixable Viability Dye eFluor 780 (FisherScientific) for 10 min, washed with FACS buffer, then fixed and permeabilized using the eBioscience Foxp3/Transcription Factor Staining Buffer Set (IThrermoFisher). Cells were incubated in 1X Fixation/Permeabilization buffer for 25 min and washed with 1X Permeabilization buffer. Prior to intracellular staining, nonspecific binding was blocked by incubation with 1% normal mouse serum and 1% normal rat serum for 10 min. Cells were then stained with fluorophore-conjugated antibodies specific for intracellular antigens. Finally, cells were washed once with 1X Permeabilization buffer, and once with FACS buffer prior to resuspension in FACS buffer for analysis on the flow cytometer.

Flow cytometry analysis and single cell sorting was done using a 4-laser 20-parameter BD FACSAria III with FACSDiva v8.0.1 software (BD Biosciences). Single cells were sorted into 384- well plates while index sorting software (BD Biosciences) simultaneous collect the phenotype data of each single cell sorted into each well. Flow cytometry data analysis and single cell index data analysis was performed using FlowJo v10 (TreeStar).

### Integrated quantitative and targeted single cell RNA-seq (qtSEQ)

Single cell transcriptional analysis was carried out using a custom RNA sequencing protocol called quantitative targeted single cell sequencing (qtSEQ). This technique sequences polyA mRNA using a nested set of 3’ targeted primers to amplify specific gene sets and is previously described ^23, 39^.

Single cells were index sorted from Foxp3^Thy1.1^ reporter mice: naïve CD4 T cells (B220-CD8- TCRb+CD4+ Thy1.1- CD44-CD62L+), T follicular helper cells (TFH, B220-CD8- TCRb+CD4+ Thy1.1- CD62L-CD44+ Cxcr5+), T regulatory cells (Treg, B220-CD8- TCRb+CD4+ Thy1.1+ Cxcr5-), T follicular regulatory cells (TFR, B220-CD8- TCRb+CD4+ Thy1.1+ CD62L- Cxcr5+) and either CD25- or CD25+. Doublets were excluded using SSC-A and SSC-H parameters. Cells were sorted from the spleens of 6 individual steady state and 6 individual mice six days after anti PD- 1 treatment (n=3 experiments) into 384-well plates at 4C.

Each well contained 1µL of reverse transcription (RT) master mix consisting of 0.03μL SuperScript II taq, 0.2μL SuperScript II 5x buffer, 0.012μL of 25μM dNTPs, 0.03μL DTT (0.1M stock), 0.03μL RNaseOUT (Invitrogen), 0.19μL DNase/RNase Free Water and 0.5μL of extended oligoDT primers (1μM stock, custom made, IDT). To allow assessment of contamination from sample processing, 8 negative control wells with no cell sorted were interspersed throughout each plate.

Immediately after sorting, reverse transcription (RT) was performed at 42°C for 50 mins then at 80°C for 10 mins. Following RT, all 384 wells were pooled within each plate and excess oligoDT primer removed using ExoSAP-IT Express (Affymetrix) at 37C for 20mins. cDNA was purified and volume reduced using a 0.8x AMpure XP SPRIselect magnetic bead (BeckmanCoulter) solution to 1x library ratio. For each plate two nested PCR reactions were used to amplify specific gene targets then a final PCR reaction performed to attach Illumina sequencing adapters. Library preparation was carried out on each plate separately to preserve cell labeling.

PCR 1 was performed in a 60μL reaction: 35μL cDNA library in H2O, 1.5μL RNaseH (5000U/mL, NEB), 1.2μL Phusion DNA polymerase (2000U/mL, NEB), 12μL 5X Phusion HF Buffer (NEB), 0.85μL of 25μM dNTPS (Invitrogen), 1.2μL of 100μM RA5-overhang primer (custom made, 5’AAGCAGTGGTGAGTTCTACAGTCCGACGATC 3’) and 25nM final concentration for each specific gene targeting external primer using the following thermocycling conditions: 37°C for 20 minutes, 98°C for 3 minutes, followed by 10 cycles of 95°C for 30 seconds, 60°C for 3 minutes, 72°C for 1 minute and final elongation at 72°C for 5 minutes. Removal of excess primers and volume reduction was performed again using 0.9x AMpure XP SPRIselect beads to 1x library ratio.

PCR2 was performed in a 20μL reaction: 10μL PCR1 reaction elute, 0.4μL Phusion DNA polymerase (2000U/mL, NEB), 4μL 5X Phusion Buffer (NEB), 0.3μL of 25μM dNTPs (Invitrogen), 2μL of 20μM RA5 (custom made, 5’ GAGTTCTACAGTCCGACGATC 3’), and 25nM final concentration for each specific gene targeting internal primer using the following thermocycling conditions: 95°C for 3 minutes followed by 10 cycles of 95°C for 30 seconds, 60°C for 3 minutes, 72°C for 1 minute and final elongation at 72°C for 5 minutes. Removal of excess primers and volume reduction was performed using AMpure XP SPRIselect beads at a 0.9x bead to 1x library ratio.

PCR3 was performed in a 20μL reaction: 10μL PCR2 reaction elute, 0.4μL Phusion DNA polymerase (2000U/mL, NEB), 4μL 5X Phusion Buffer (NEB), 0.3μL of 25μM dNTPs (Invitrogen), 2μL of 10μM RP1 primer (Illumina), 2μL of 10μM RPI library specific primer (Illumina) and 1.3μL water using the following thermocycling conditions: 95oC for 3 minutes followed by 8 cycles of 95°C for 15 seconds, 60°C for 30 seconds, 72°C for 30 seconds and final elongation at 72°C for 5 minutes. Removal of excess primers and volume reduction was performed using 0.7x AMpure XP SPRIselect beads to 1x library ratio and then eluted in 20μL H2O for quantification and library sequencing preparation.

Final cDNA amplified library concentration and quantity was determined using a Qubit 4 Fluorometer (Invitrogen Technologies) and 2100 BioAnalyzer (Agilent). Separately labeled cDNA libraries were mutliplexed according to manufacturer’s protocol (Illumina, San Diego CA). Libraries were sequenced using an Illumina NextSeq 500 (Read 1: 19 cycles; Index Read 1: 6 cycles; Read 2: 67 cycles).

### Data processing and normalization

Bcl2fastq (Illumina) was used to demultiplex sequencing libraries using plate barcodes and assort reads by well barcodes prior to sequence alignment using Bowtie2 (v2.2.9) to a custom genome consisting of the amplicon regions for the specifically targeted genes used in the experiment based on the murine genome GRCm38.p6, version R97 (Ensembl). HTseq was used for read and UMI reduced tabulation. Custom scripts for these steps available upon request.

Quality control included T cells that contained at least one count of *Cd3d, Cd3e, or Cd3g and* excluded cells that contained fewer than 60 UMI or more than 2000 counts and had fewer than 25 unique genes detected. A total of 2272 CD4 T cells from 3 experiments (n=6 steady state and n=6 anti PD-1 treat mice) were analyzed.

Data from all mice and experiments was pooled into a single count matrix for processing using the R package Seurat (v3.2.3) ^80^; scaling and normalization was carried out separately on each sorted cell phenotype using the ScTransform package (v0.3.2.9002) ^81^ with the batch_var option set to the metadata variable “Experiment”. This normalized UMI count matrix was then used for downstream analysis.

### scRNA Sequencing Computational Analysis

For differential gene expression analyses, MAST (Model-based Analysis of Single Cell Transcriptomics) package (v1.16.0 https://github.com/RGLab/MAST/ ) ^82^ using a hurdle model was used to generated fold change and p-values for all genes. Volcano plots were then created using Prism v9.0 (GraphPad Software). The R package pheatmap (v1.0.12) was used to plot heatmaps of averaged UMI counts with euclidean distance hierarchical clustering. Single cell heatmaps were generated using the “Doheatmap” function and dot plots using the “DotPlot” function from the Seurat package. Top significant marker genes were identified between clusters using the FindMarkers function in Seurat. Uniform Manifold Approximation and Projection (UMAP) was performed using Seurat.

Network analysis was performed using the web application Metascape ^44^. Multiple lists of significantly upregulated genes from TFH1 and TFH2 cell subsets as compared to naive T cells were inputed into the program. Species analyzed was chosen to be human due to the superior database of protein-protein interactions. All other default settings were used including the molecular complex detection algorithm (MCODE) which highlights densely connected networks that are more likely to be enriched in functional pathways. Predicted ligand receptor interactions were calculated and visualized using Cellphone DB ^52^ and Connectome ^50^. Statistical tests were calculated and graphed using Prism v9.0 (GraphPad Software). A p-value *<0.05, **<0.01,

***<0.001 was considered statistically significant.

## ACKNOWLEDGEMENTS

This work was supported by the US National Institutes of Health (AI047231, AI040215 and AI071182) and Bill & Melinda Gates Foundation (BMGF 0PP1154835) to M.G.M.-W.

## AUTHOR CONTRIBUTIONS

A.G.S., B.W.H., K.B.M., L.M.W., M.M.W. designed and performed experiments, A.G.S., B.W.H., L.M.W., M.M.W. developed single cell qtSEQ, A.G.S., L.M.W., M.M.W. designed the study, analyzed data and wrote the paper.

## Extended Data Figures

**Extended Data Fig 1.**
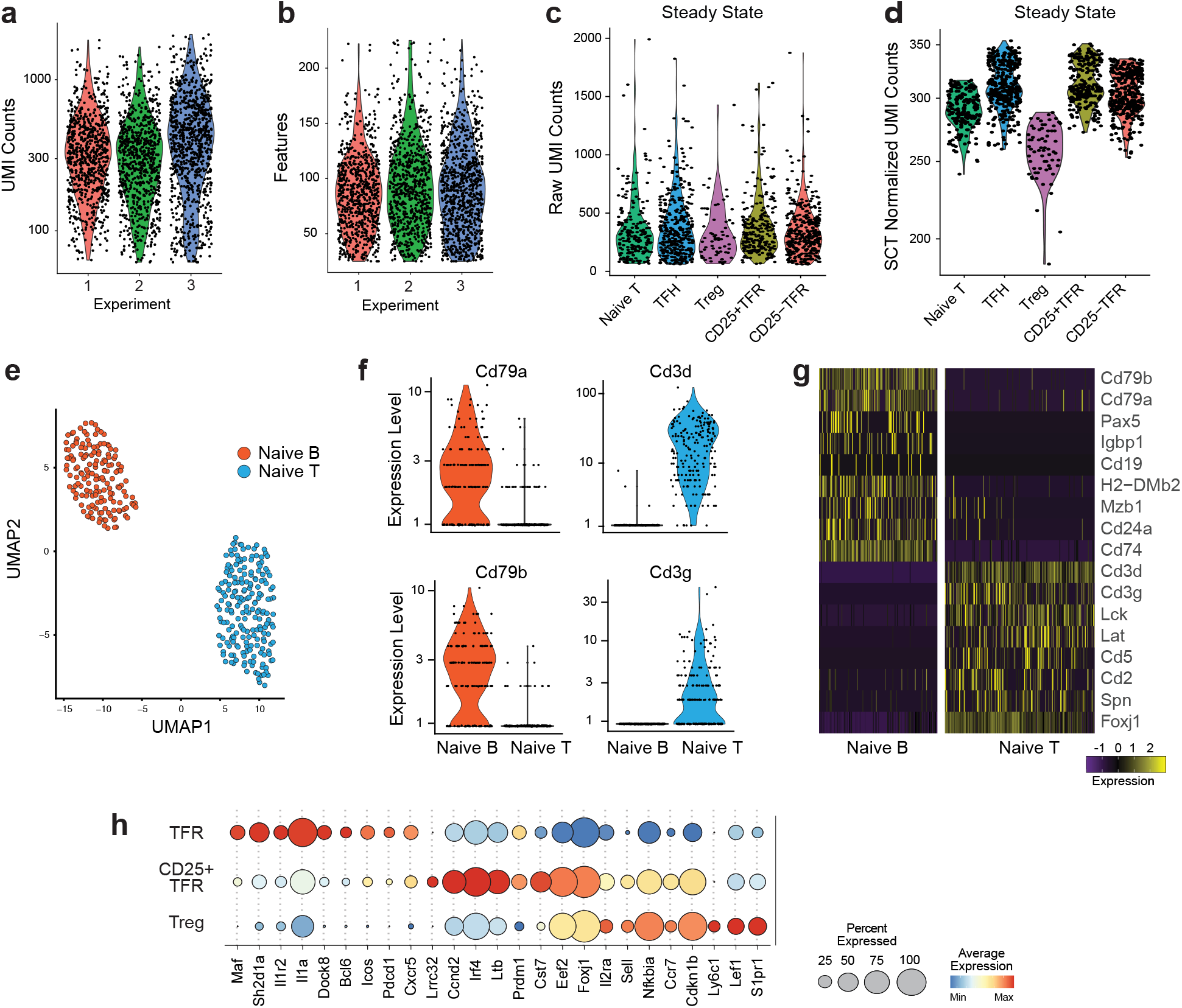
Quantitative and targeted single cell RNA-sequencing (qtSEQ) of splenic CD4 T cells. **a)** Single cell RNA-seq (qtSEQ) violin plot of UMI counts displayed on a log scale for individual experiments. Each black dot represents one cell. Experiment 1 n=727 cells, experiment 2 n=752, and experiment 3 n=879. **b)** Number of features (gene products) detected for each cell per experiment. **c)** Steady State raw pre-normalized UMI counts per cell for each sorted subset of CD4 cell and **d)** after post normalization using scTransform. Cells with UMI counts of <60 or >2000 were filtered out. **e)** UMAP unbiased clustering of gene products for naive B cells (Gr1-FceR1a- IgD+IgMlow CD138-GL7- CD79b+CD19+CD38+) and naïve CD4 T cells (B220-CD8- TCRb+ CD4+Thy1.1- CD44- CD62L+) n=200. **f)** Violin plots with single cell overlay of B cell gene products Cd79a and Cd79b (left) and T cell gene products Cd3d and Cd3g (lright) expression levels for naïve B (red) and naïve T (blue) cells. **g)** Single cell heatmap of top significant markers gene products for naïve B and T cells. **h)** Dot plots displaying average gene expression of sorted cells for TFR CD25-, TFR CD25+ and Treg cells. Dot size depicts the percent of cells detecting the gene product and color is average expression.

**Extended Data Fig 2.**
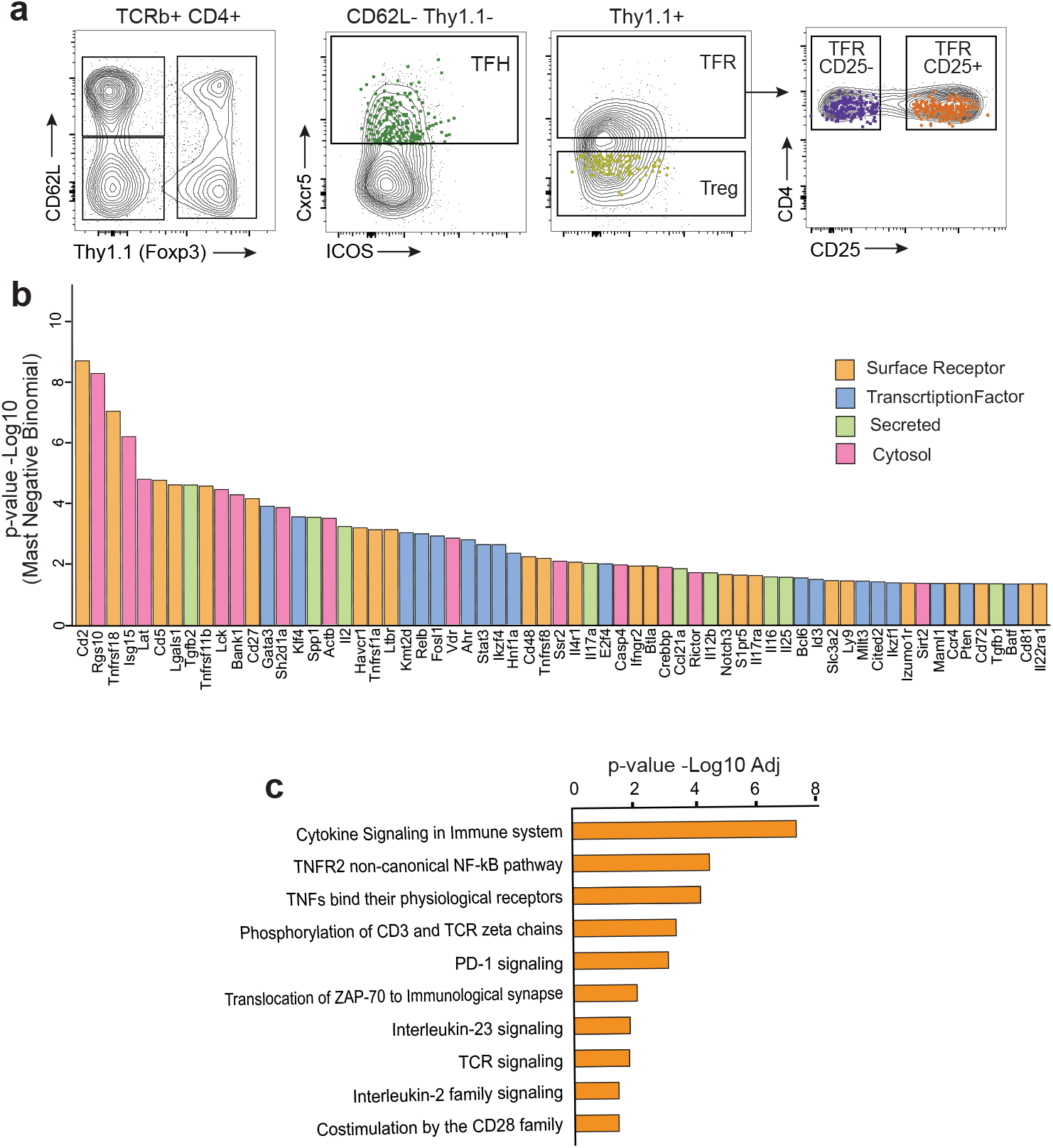
PD-1 blockade augments TFH program. **a)** Flow cytometric analysis of Foxp3Thy1.1 reporter mice TCRb+ CD4+ T cells from anti PD-1 treated mice with overlaid representative index sorted cells. TFH (Thy1.1- CD44+ CD62L- Cxcr5+) n=418 green, Treg (Thy1.1+ Cxcr5-) n=82 yellow, TFR (Thy1.1+ Cxcr5+) either CD25- n=379 purple, or CD25+ n=253 orange. **b)** Top significant differentially expressed gene products from TFH cells from steady state versus anti-PD1 treated mice. Gene products are ranked by descending order of significance (p-value -Log10 Mast Negative Binomial) and color coded according to cellular localization as either surface receptor (orange), transcription factor (blue), secreted (green) and cytosol (pink). **c)** Selected significantly enriched pathways against the Reactome gene set for TFH cells from steady state and anti PD-1 treated mice plotted by descending p-value.

**Extended Data Fig 3.**
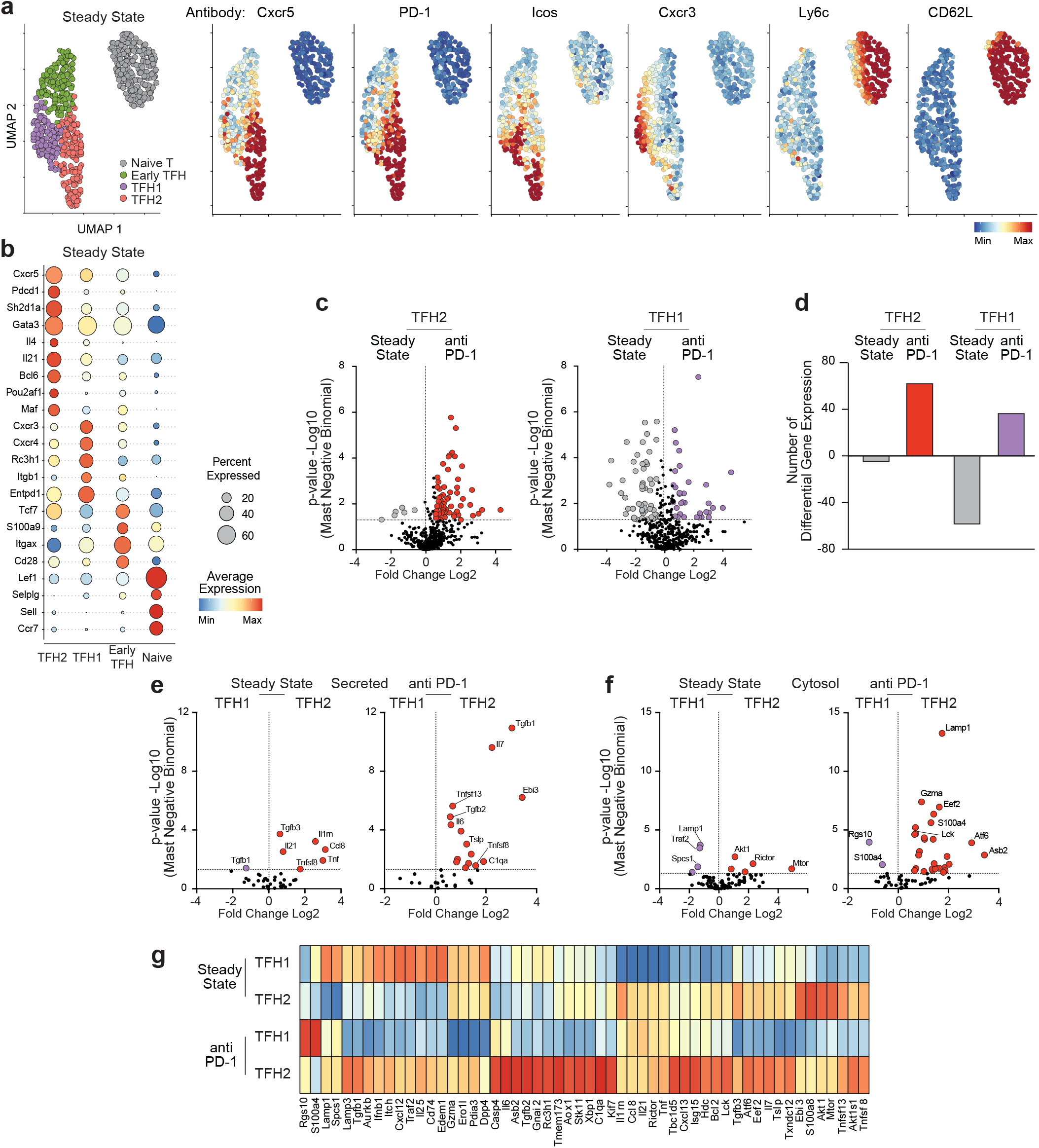
Differential impact of PD-1 blockade on subsets of TFH. **a)** Foxp3Thy1.1 reporter mice UMAP flow cytometry analysis of sorted naïve T cells and TFH cells from steady state mice. Six parameters (Cxcr5, PD-1, Icos, Cxcr3, Ly6c, CD62L) were used to cluster the cells (left). The clusters identified naïve T cells (grey), early TFH (green), TFH1 (purple) and TFH2 (salmon) cells. Distribution of individual antibodies Cxcr5, PD-1, Icos, Cxcr3, Ly6c, and CD62L are overlaid with each fluorophore (right panels) intensity from minimum (blue) to maximum (red). **b)** Dot plots displaying average gene expression of sorted cells for TFH cells from steady state mice clustered as TFH2, TFH1, early TFH (defined from unbiased UMAP projections in Fig 3a-d) and naïve T cells. Most significant marker genes are shown with dot size depicting the percent of cells detecting the gene product and color is the average expression. **c)** Volcano plots comparing TFH2 (left) and TFH1 (right) from steady state and anti PD-1 treated mice. Large and colored dots were considered significant (>1.5 Log2 fold change and p<0.05). **d)** Changes in the number of differential gene expression products from c) visualized in a bar graph. **e)** Volcano plots comparing secreted and **f)** the cytosol gene products from the TFH1 cluster (n=84) and TFH2 cluster (n=129) at steady state or in the same clusters after PD-1 blockade (anti PD-1: TFH1 cluster n=167, TFH2 n=228). Large and colored dots were considered significant (>1.5 Log2 fold change and p<0.05). **g)** Selected significant gene products (>1.5 Log2 fold change and p<0.05) from e) and f) displayed is an averaged single cell heatmap.

**Extended Data Fig 4.**
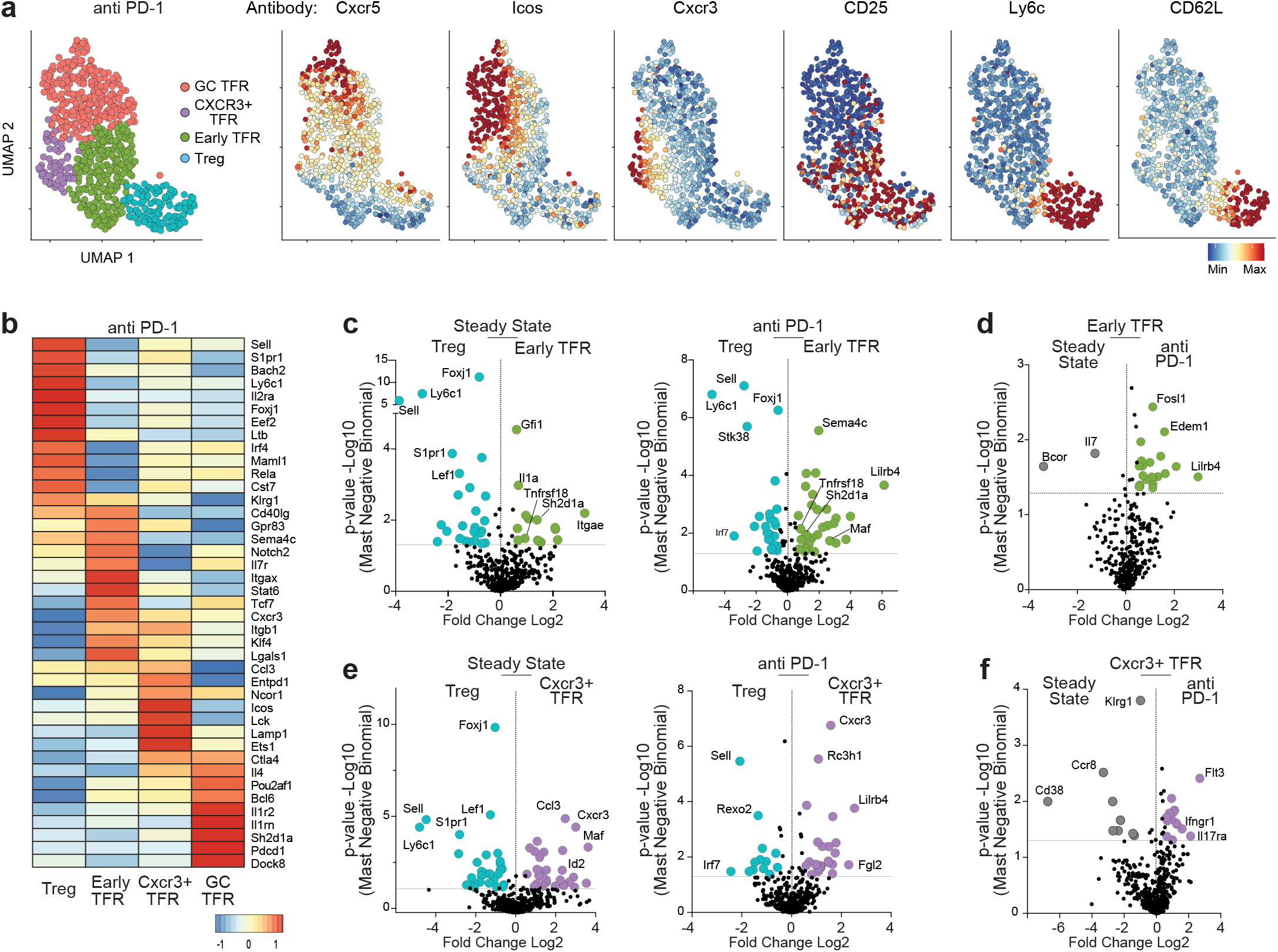
PD-1 blockade of TFR cell subsets. **a)** Foxp3Thy1.1 reporter mice UMAP flow cytometry analysis of anti PD-1 sorted CD4 T (CD4+ TCRb+) cells, regulatory Treg (Thy1.1+ Cxcr5-), TFR CD25- (Thy1.1+, Cxcr5+, CD25-) and TFR CD25+ (Thy1.1+, Cxcr5+, CD25+) cells. Five parameters (Cxcr5, Icos, Cxcr3, Ly6c, CD62L) were used to cluster the cells (left). The 4 Foxp3 (Thy1.1+) clusters identified Treg (colored blue), early TFR (green), Cxcr3+ TFR (purple) and GC TFR (salmon) cells. Distribution of individual antibodies Cxcr5, Icos, Cxcr3, CD25, Ly6c, CD62L are overlaid with each fluorophore intensity (right panels) from minimum (blue) to maximum (red). **b)** Single cell qtSEQ averaged heatmap of the top significant marker genes differentiating each regulatory cell cluster from anti PD-1 treated mice (n=714). **c)** Gene expression volcano plots of Treg compared to early TFR cells at steady state (left) and anti PD-1 Treg to early TFR cells (right). Displayed are Log2 fold change and Mast negative binomial -Log10 p-value. Large and colored dots are gene products that have >1.5 Log2 fold change and/or a p<0.05. **d)** Volcano plot of the gene products from early TFR cells from steady state and anti PD-1 treated mice. **e)** Volcano plots of Treg compared to Cxcr3+ TFR cells from steady state (left) and anti PD-1 treated mice (right). **f)** Volcano plot of the gene products from Cxcr3+ TFR cells from steady state and anti PD-1 treated mice.

**Extended Data Fig 5.**
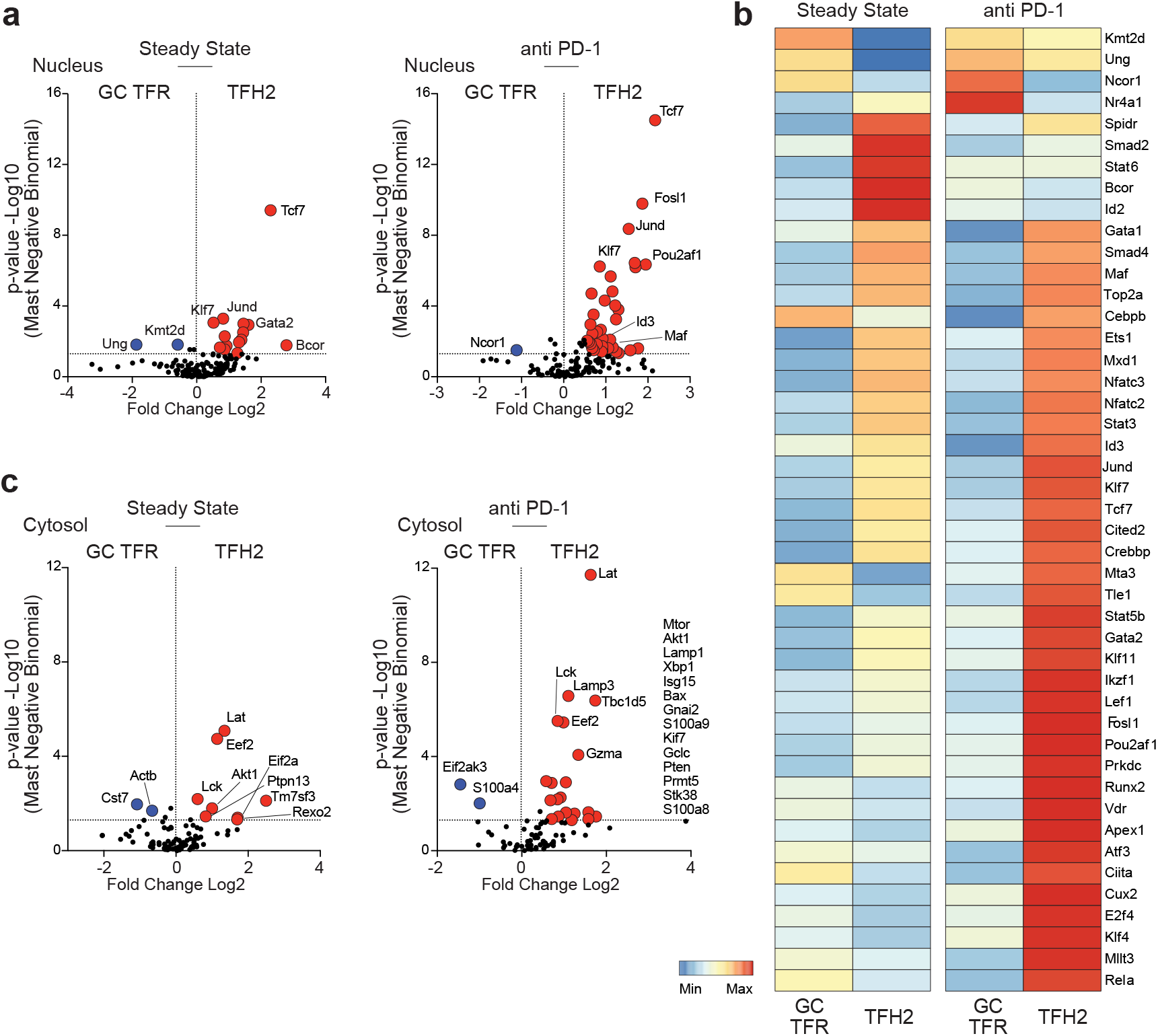
PD-1 blockade alters GC regulatory dynamics. **a)** Single cell gene expression volcano plots comparing genes encoding proteins localized in the nucleus from steady state GC TFR (n=221) and TFH2 cells (n=129) (left) and after PD-1 blockade (GC TFR n=272, TFH2 n=228 cells) (right). Large and colored dots were considered significant (>1.5 Log2 fold change and p<0.05). **b)** Averaged single cell heatmap of the significant cell nucleus products identified in a). **c)** Volcano plots comparing steady state and anti PD-1 treated GC TFR and TFH2 cells from cytosolic gene products.

**Extended Data Fig 6.**
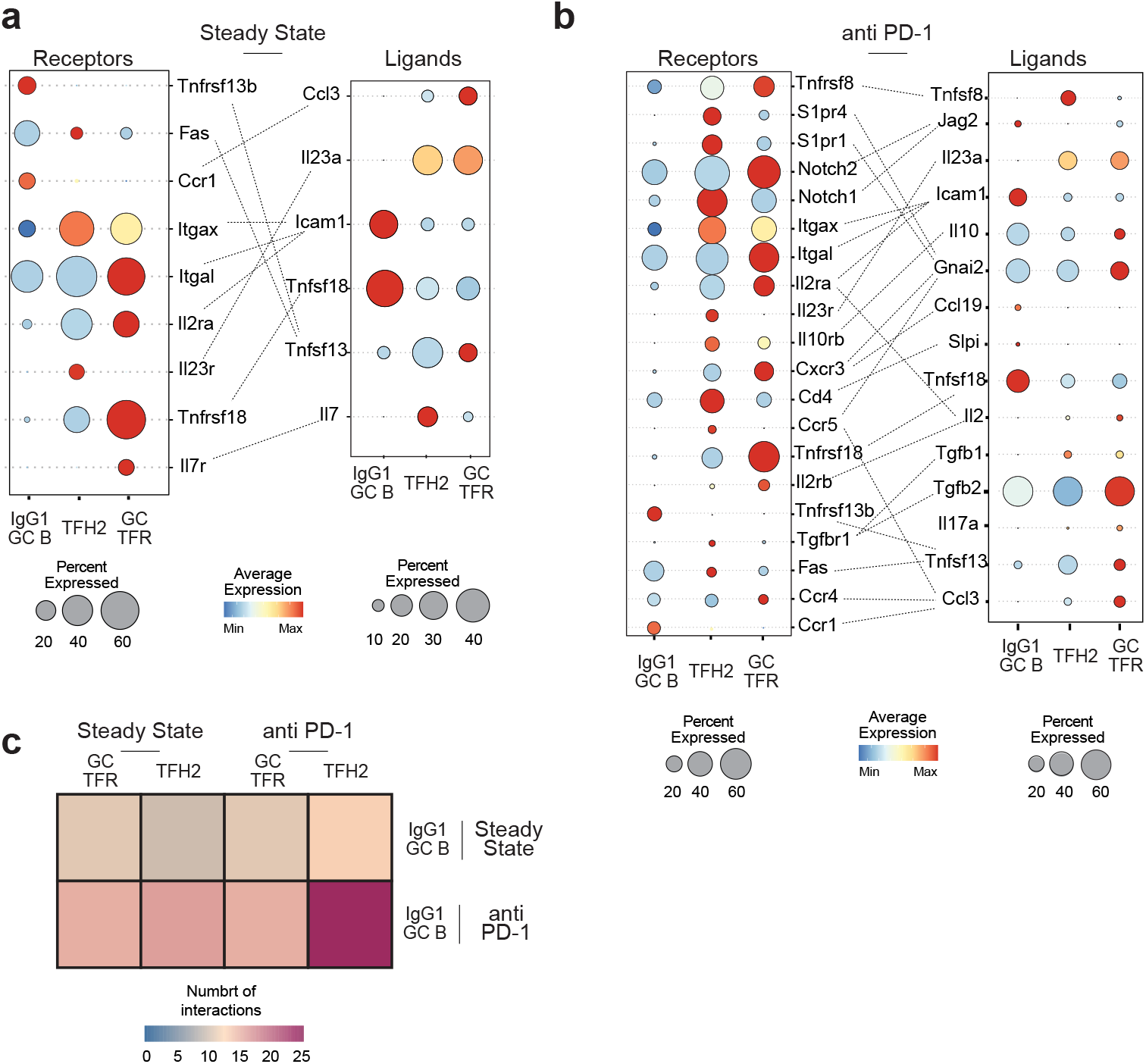
Ligand and receptor interactions for steady state and PD-1 blockade. **a)** Single cell Connectome cell-cell connectivity patterns by ligand-receptor mapping of IgG1 GC B, GC TFR, and TFH2 cells from steady state and **b)** anti PD-1 treated mice. Dot plots link to the Circos / network plots in Fig 7a. Connections are filtered by significance (p<0.05) and minimum fraction of cells expressing a gene product (>10%). Dot size depicts the percent of cells detecting the gene product and color gives average expression. **c)** Heatmap of the total number of significant (p<0.05) interactions between GC TFR and TFH2 T cells and IgG1 GC B cells from steady state and anti PD-1 treated mice. Links to the CellPhoneDB receptor-ligand interactions in Fig 7b.

